# Bradycardic mice undergo effective heart rate improvement after specific homing to the sino-atrial node and differentiation of adult muscle derived stem cells

**DOI:** 10.1101/393512

**Authors:** Pietro Mesirca, Daria Mamaeva, Isabelle Bidaud, Matthias Baudot, Romain Davaze, Mattia L. DiFrancesco, Violeta Mitutsova, Angelo G. Torrente, Nikola Arsic, Joël Nargeot, Jörg Striessnig, Amy Lee, Ned J. Lamb, Matteo E. Mangoni, Anne Fernandez

**Author notes:** These authors contributed equally to this work (Pietro Mesirca, Daria Mamaeva).

## Abstract

Current treatments for heart automaticity disorders still lack a safe and efficient source of stem cells to restore normal biological pacemaking. Since adult <u>M</u>uscle-<u>D</u>erived <u>S</u>tem <u>C</u>ells (MDSC) show multi-lineage differentiation *in vitro* including into spontaneously beating cardiomyocytes, we questioned whether they could effectively differentiate into cardiac pacemakers, a specific population of cardiomyocytes producing electrical impulses in the sino-atrial node (SAN) of adult heart. We show here that beating cardiomyocytes, differentiated from MDSC *in vitro*, exhibit typical characteristics of cardiac pacemakers: expression of markers of the SAN lineage Hcn4, Tbx3 and Islet1, as well as spontaneous calcium transients and hyperpolarization-activated “funny” current and L-type Ca_v_1.3 channels. Pacemaker-like myocytes differentiated *in vitro* from Ca_v_1.3-deficient mouse stem cells produced slower rate of spontaneous Ca^2+^ transients, consistent with the reduced activity of native pacemakers in mutant mice. *In vivo*, undifferentiated wild type MDSC migrated and homed with increased engraftment to the SAN of bradycardic mutant Ca_v_1.3^-/-^ within 2-3 days after systemic I.P. injection. The increased homing of MDSCs corresponded to increased levels of the chemokine SDF1 and its receptor CXCR4 in mutant SAN tissue and was ensued by differentiation of MDSCs into Ca_v_1.3-expressing pacemaker-like myocytes within 10 days and a significant improvement of the heart rate maintained for up to 40 days. Optical mapping and immunofluorescence analyses performed after 40 days on SAN tissue from transplanted wild type and mutant mice showed MDSCs integrated as pacemaking cells both electrically and functionally within recipient mouse SAN. These findings identify MDSCs as directly transplantable stem cells that efficiently home, differentiate and improve heart rhythm in mouse models of congenital bradycardia.

## Introduction

Stem cells with the capacity to self-renew and differentiate into several lineages can be isolated from most adult tissues and organs. However, as the largest vascularised and innervated organ in the body, and one with the life-long ability to undergo atrophy and regeneration, skeletal muscle is particularly rich in multipotent stem cells (reviewed ^1,2,3^). Various populations of stem cells have been isolated from muscle, the best studied being satellite cells. Named from their tight association to muscle fibres, satellite cells account for muscle fibre regeneration and are now recognized as a heterogeneous population (reviewed ^4^). In addition, skeletal muscle contains connective tissue-, vascularisation-, and innervation-derived progenitor and stem cells, which cooperate with satellite cells in tissue repair (reviewed ^1,3,5^). Consistent with the neuro-muscular identity of skeletal muscle, differentiation of muscle-derived stem cells into different meso-ectodermal cell lineages was reported in several studies including from human muscle ^6-8^ and after clonal growth of muscle stem cells ^9^.

We previously reported that a population of <u>m</u>uscle-<u>d</u>erived <u>s</u>tem <u>c</u>ells (MDSCs) ^10^ isolated from adult mouse muscle on the basis of their low adherence by sequential preplating ^11,12^ is multipotent ^10^. In particular this population spontaneously differentiated into autonomously beating cardiomyocyte-like cells ^10^. This sustained autonomous phenotype questioned whether beating cardiomyocytes derived from MDSC may be similar to pacemaker myocytes, a small population of highly specialized cells in the right atrium of adult hearts located in the Sino-Atrial (SAN) and Atrio-Ventricular Node (AVN)^13^. Pacemaker cells are endowed with automatic electrical activity, and this enables them to generate the cardiac pacemaker action potential. This is conducted through the atrial tissue and the AVN to the Purkinje fibre network from which it depolarizes the ventricular myocardium^13^. In vertebrates, automaticity is one of the first features of cardiogenesis, with all cardiac cells of the early embryo capable of autonomous beating. During late development of the chambers and conduction system, the SAN emerges from a morphologically distinct area that develops within the sinus venosus, at the site where the right atrium enters the intercaval region, due to up-regulation of the Tbx18 and Tbx3 transcription factors and their repression of cardiogenic Nkx2.5^14,15^. The generation and conduction of the pacemaker impulse within the SAN differs from that in working atrial myocytes in that membrane depolarisation is spontaneous and intercellular electrical resistance is high ^16,17^. To fulfil this requirement, pacemaker myocytes uniquely express a particular set of ion channels: the hyperpolarisation-activated cAMP-gated cation channel 4, HCN4; voltage-gated Ca_v_1.3 channels; and a specific subset of connexins, including connexin45 (Cx45) and Cx30.2 (but not the high-conductance gap junction proteins Cx40 and Cx43 which are typically expressed in working cardiomyocytes) ^18,19^. Pacemaker myocytes thus represent a specialized type of cardiac myocyte that is essential for proper heart function and whose properties and defining markers are distinct from those of working cardiomyocytes.

In humans, SAN dysfunction results in reduced life expectancy and necessitates the implantation of >450,000 pacemakers/year in Europe and the U.S., a number predicted to double over the next half century ^20^. Whereas pacemaker implantation remains the primary therapy for chronic SAN dysfunction and bradycardia-associated arrhythmias ^21^, this approach is costly, requiring lifelong management including regular technical inspections. These limitations and complexity in the use of an electronic device are even greater in pediatric and young patients ^22^. Considering the global impact of SAN dysfunction, it is generally agreed that new therapeutic strategies are needed. The development of a cell-therapy approach using patient-derived cells to repair the damaged SAN and restore pacemaker activity appears as a novel promising strategy to achieve stable, life-long improved heart function and to restore patient quality of life ^22^. Toward this goal it will be necessary to identify suitable sources of transplantable stem cells that are capable of differentiating *in vivo* into myocytes with functional pacemaker activity.

In the current study, we tested the ability of MDSCs to differentiate, both *in vitro* and *in vivo* in mutant mice, into pacemaker myocytes. We report that MDSC can migrate and home to the SAN where they differentiate into pacemaker-like cardiomyocytes. The resulting functional integration of MDSCs significantly ameliorates heart rhythm in congenitally bradycardic mutant mice.

## Results

### *In vitro* analysis: MDSCs spontaneously differentiate into autonomously beating cardiomyocytes with the characteristics of pacemaker cells

MDSCs were isolated from mouse hind-limb muscle via sequential preplating of non-adherent cells, as shown in Fig. 1a and described previously ^10^. Non-adherent stem cells were replated in fresh proliferation medium every 24 hours to avoid their premature differentiation and to stimulate their proliferation. The rationale for daily replating and centrifugation is two-fold: daily replating prevents attachment and loss of non-adherent stem cells, which would otherwise attach to rapidly adhering and proliferating cells (e.g., fibroblasts, myofibroblasts and pre-adipocytes); and daily centrifugation depletes the cultures of muscle fibre-derived microsomes and other muscle-specific non-cellular components that may contribute to the myogenic commitment of muscle stem cells. A comparative analysis of the transcriptomes of muscle cells extracted 24 hours after activation of proliferation (corresponding to PP1) and MDSCs isolated after 7 days of pre-plating (corresponding to PP8) revealed a 12-fold increase in cyclinA2 expression (Fig.1b), indicating an increased level of cell proliferation. The same analysis revealed a 10-fold increase in the expression of Islet1, a LIM-homeodomain transcription factor expressed in heart and pancreatic progenitor/stem cells, and a >30-fold increase in expression of Peg3/PW1, which is expressed in stem cells isolated from muscle and several adult tissues ^23^. Of note, the stemness factor Nestin, whose expression is high in PP1 cells, was increased >2-fold in PP8 MDSCs (Fig. 1b). These data confirm that 7 days of preplating non-adherent cells from mouse skeletal muscle is an effective means of isolating a proliferating stem cell population.

**Fig 1.**
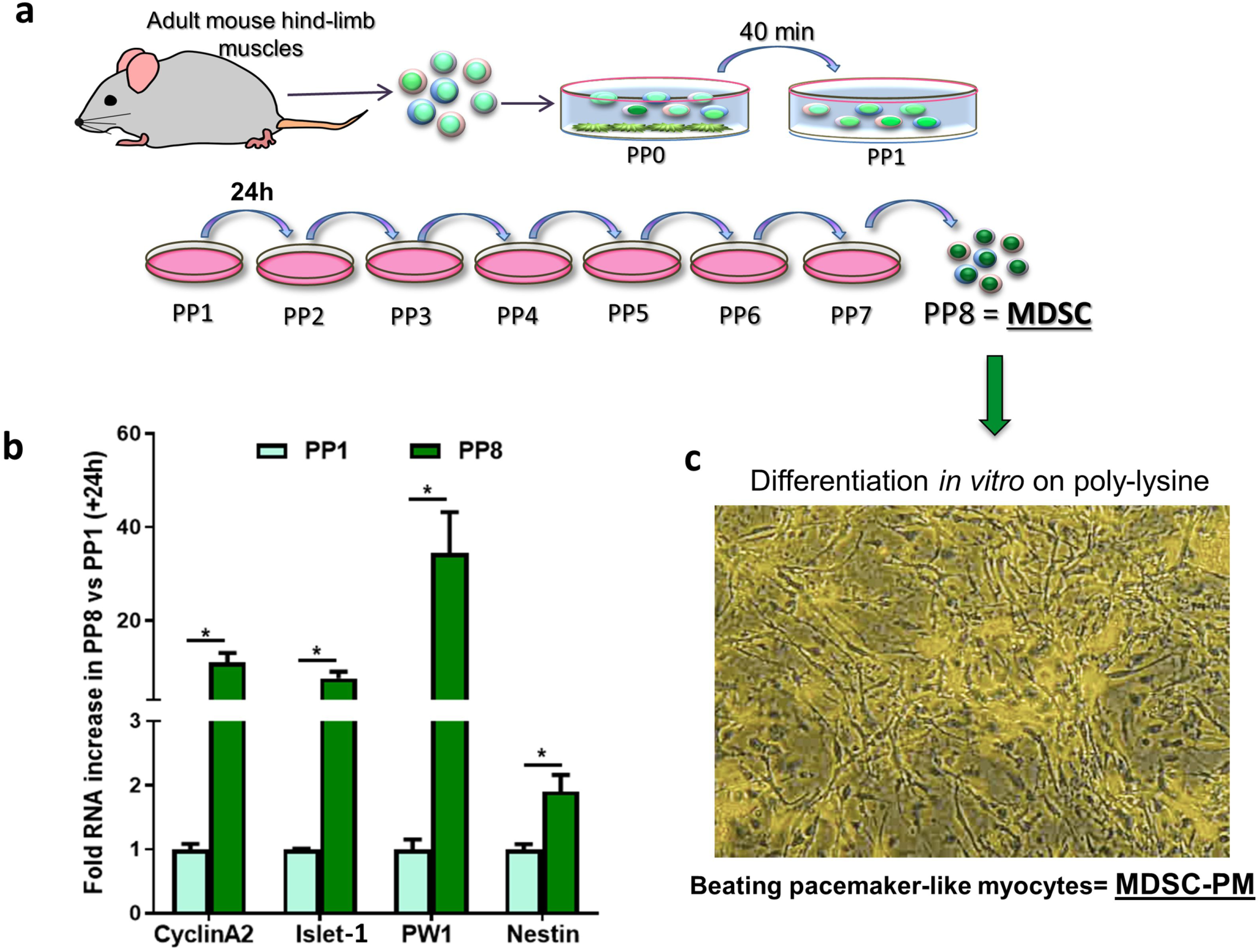
Muscle stem cell isolation, expression of stemness factors, and differentiation of pacemaker-like cells *in vitro*. (**a**) Schematic overview of the protocol for isolating non-adherent muscle-derived stem cells (MDSCs) from mouse hind-limb muscle. (**b**) Levels of RNA expression for *CyclinA2, Islet1, Peg3/PW1*, and *Nestin* in PP1 muscle cells cultured for 24 hours versus MDSC PP8 cells, as determined by RNAseq profiling of independent replicates. Shown are the fold increase in PP8 MDSCs vs. PP1 cells (levels in PP1 cells normalized as 1x). Statistical significance was tested for each gene using the unpaired t-test. *p<0.05. Error bars indicate s.e.m. (**c**) Phase-contrast image of MDSCs differentiated into <u>m</u>uscle <u>s</u>tem <u>c</u>ell-derived <u>p</u>ace<u>m</u>aker myocytes (MDSC-PMs) after 3 weeks on poly-lysine. The field is the same as that shown in Supplementary video 1.

We previously reported that non-adherent MDSCs in PP8 cultures provided with an attachment matrix such as laminin, fibronectin, or a fibroblast layer (but not collagen) became adherent and underwent multi-lineage differentiation^10^. Amongst the different phenotypes observed, cells with the ability to beat autonomously were observed within 3 to 5 days and these cells exhibited persistent contractile activity for over 2 months in culture (Fig.1c and corresponding Supplementary video 1). Individually beating cells or clusters of cells with coordinated beating rhythms were observed (Supplementary videos 2 and 3), and the morphology of these beating myocytes was reminiscent of that of native SAN pacemaker cells and not that of atrial cardiomyocytes (Supplementary Fig. 1a-d). Henceforth, we refer to these *in vitro* differentiated, spontaneously beating cells as MDSC<u>-</u>derived <u>p</u>ace<u>m</u>aker-like cells (MDSC-PMs). To investigate whether these cells exhibit properties of SAN cardiomyocytes, we first analyzed a number of molecular markers of pacemaker development and activity. SAN development is initiated in a small area of the septum venosus that co-expresses Tbx18 and Islet1 transcription factors, after which expression of the Shox2 and Tbx3 transcription factors is initiated and that of Nkx2.5 is down-regulated ^14,24,25^. Initial RT-PCR analysis of non-differentiated MDSCs showed RNA expression of Tbx18, Shox2 and Tbx3 and no expression of Nkx2.5 (Supplementary Figure 1e, Figure 2 and supplementary Table 1). These results were confirmed and extended by comparative RNAseq analysis carried out on non-differentiated MDSCs, MDSC-PMs differentiated into pacemaker-like cells and native right atrium (RA) and sinoatrial (SAN) tissue sample (Figure 2a and Figure 3). Notably, results shown Figure 2a show the expression of Islet1 in differentiated MDSC-PM. In addition, expression of Shox2, a core component in the development of the venous pole and pacemaker^25^ and in pacemaker program in embryoid bodies^26^, is markedly increased in differentiated MSDC-PM to a level similar to the expression measured in native SAN plus RA tissue (SAN-RA). Pacemaker-like differentiated MDSC-PMs likewise expressed increased levels of Tbx18, Tbx3 and Islet1 and showed no expression of Nkx2.5 or Tbx2, two transcription factors shown to be implicated in the atrial differentiation pathway^25^. This specific expression pattern is consistent with the identification of beating myocytes differentiated *in vitro* from MDSCs as being similar to cardiac pacemaker myocytes and not simply fetal-derived immature cardiomyocytes. Indeed, pacemaker cells in the adult heart differ from foetal and neonate cardiomyocytes and from adult atrial and ventricular cardiomyocytes in that they do not express Nkx2.5, which acts as a repressor of the pacemaker gene expression program ^14,25^. In addition, pacemaker myocytes are unique in retaining Islet1 expression in adult heart ^27^.

**Table 1.** Telemetric ECG intervals for transplanted Ca_v_1.3^-/-^ and wild-type mice during 12 hours resting daytime. Shown are the values of the ECG parameters and the presence of sinus (SAN) pauses and 2^nd^ –degree atrioventricular blocks (AVBII), (mean ± S.E.M.).

| | $\text{Ca}_v1.3^{-/-}$<br>before<br>injection | $\text{Ca}_v1.3^{-/-}$<br>40 days<br>post-injection | n | p value<br>$\text{Ca}_v1.3^{-/-}$ | Wild-type<br>before<br>injection | Wild-type<br>40 days<br>post-injection | n | p value<br>WT |
| --- | --- | --- | --- | --- | --- | --- | --- | --- |
| Heart rate<br>(bpm) | 310 ± 15 | 410 ± 19 | 6 | 0.012 | 568 ± 6 | 548 ± 2 | 4 | ns |
| PP (ms) | 215 ± 6 | 169 ± 6 | 6 | 0.006 | 115 ± 6 | 112 ± 1 | 4 | ns |
| PR (ms) | 55 ± 2 | 51 ± 1 | 6 | ns | 34 ± 1 | 35 ± 1 | 4 | ns |
| QRS (ms) | 17 ± 1 | 16 ± 1 | 6 | ns | 10 ± 1 | 10 ± 1 | 4 | ns |
| QT (ms) | 71 ± 5 | 66 ± 3 | 6 | ns | 43 ± 1 | 41 ± 1 | 4 | ns |
| QTc (ms) | 50 ± 3 | 52 ± 2 | 6 | ns | 41 ± 1 | 39 ± 1 | 4 | ns |
| SAN<br>pauses/60s | 0.9 ± 0.5 | 0.3 ± 0.1 | 6 | ns | 0 | 0 | 4 | ns |
| AVBII/60s | 0.5 ± 0.2 | 0.3 ± 0.1 | 6 | ns | 0 | 0 | 4 | ns |

**Fig 2.**
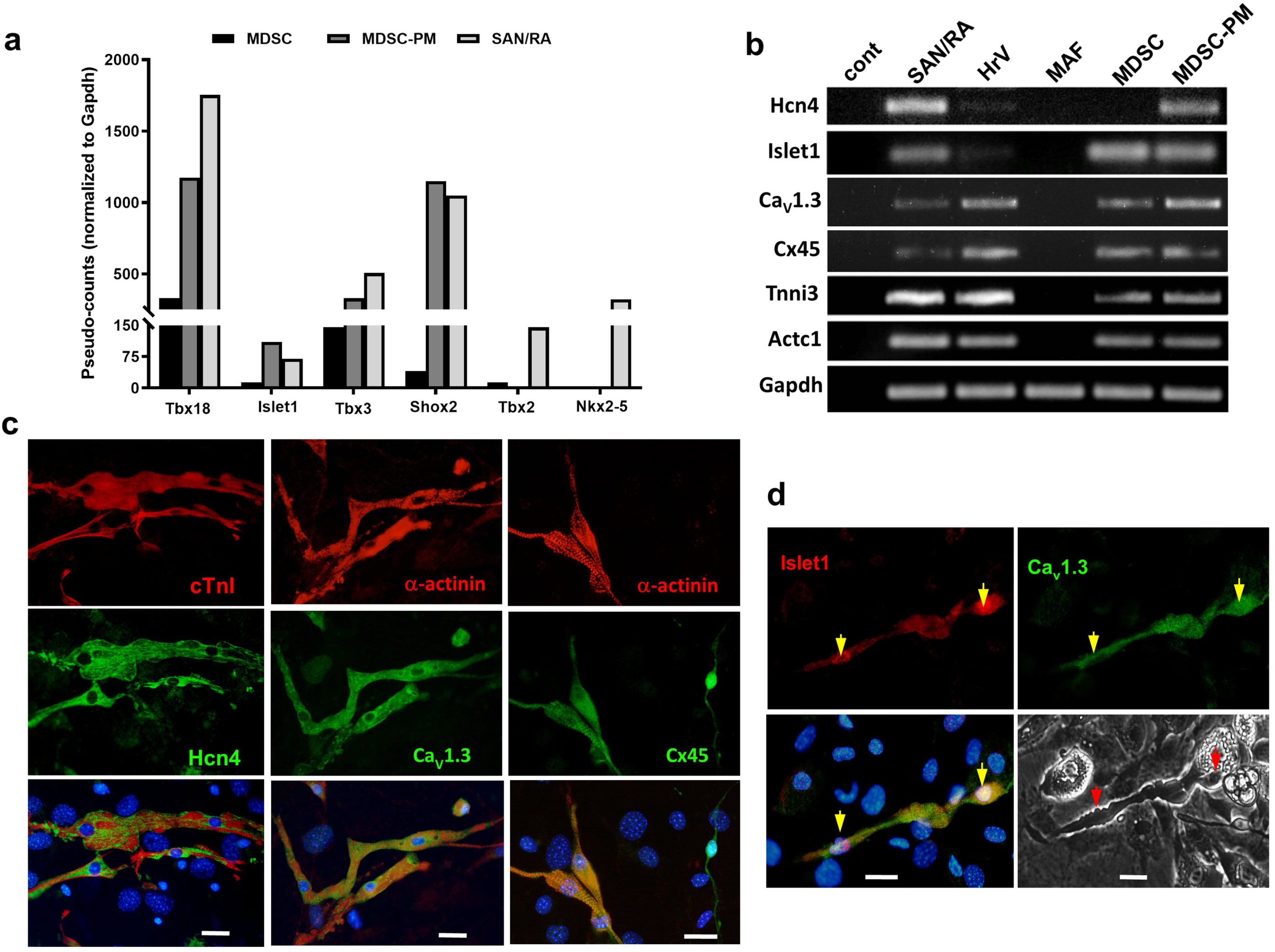
Expression of sino-atrial markers in MDSCs and in MDSC-PMs differentiated *in vitro* from MDSCs. (**a**) RNAseq analysis of the expression of transcription factors involved in sino-atrial node (SAN) development: Tbx18, Islet1, Tbx3, Shox2, Tbx2 and Nkx2.5). SAN/RA: mouse sino-atrial node and Right Atrium tissue; MDSCs: muscle-derived stem cell; MDSC-PMs: MDSCs differentiated *in vitro* into pacemaker-like beating cells. Results are shown as pseudo-counts for each gene expression, normalized in ratio to the expression level of Gadph. (**b**) RT-PCR-based analysis of expression of cardiac pacemaker-specific markers in mouse. HCN4; Islet1; Ca_V_1.3; Cx45 (Connexin45); Tnni: (cardiac Troponin I); Actc: (cardiac α-actinin). Mouse SinoAtrial Node and Right Atrium tissue (SAN/RA); muscle-derived stem cell (MDSC); MDSCs differentiated *in vitro* into pacemaker-like beating cells (MDSC-PM); heart ventricles (H(V)); mouse adult fibroblasts (MAF). Control without addition of cDNA are indicated as Cont, Gapdh was used as loading control. Results are representative of analysis of n=3 independent MDSC and MDSC-PM RNA preparations. (**c**) Co-immunostaining of MDSC-PMs for HCN4 and cardiac TroponinI (cTnI): left panels; sarcomeric α-actinin and Ca_v_1.3: middle panels; and sarcomeric α-actinin and Connexin45 (Cx45): right panels. (**d**) Co-immunostaining for Islet1 and Ca_v_1.3, and corresponding phase-contrast image of 2 beating MDSC-PMs (indicated by arrows) that are also shown live in Supplementary video S4. Immunostaining is representative of n=20 independent cultures of MDSC-PMs from wild-type mouse muscle. Scale bars, 10 µm

**Fig. 3.**
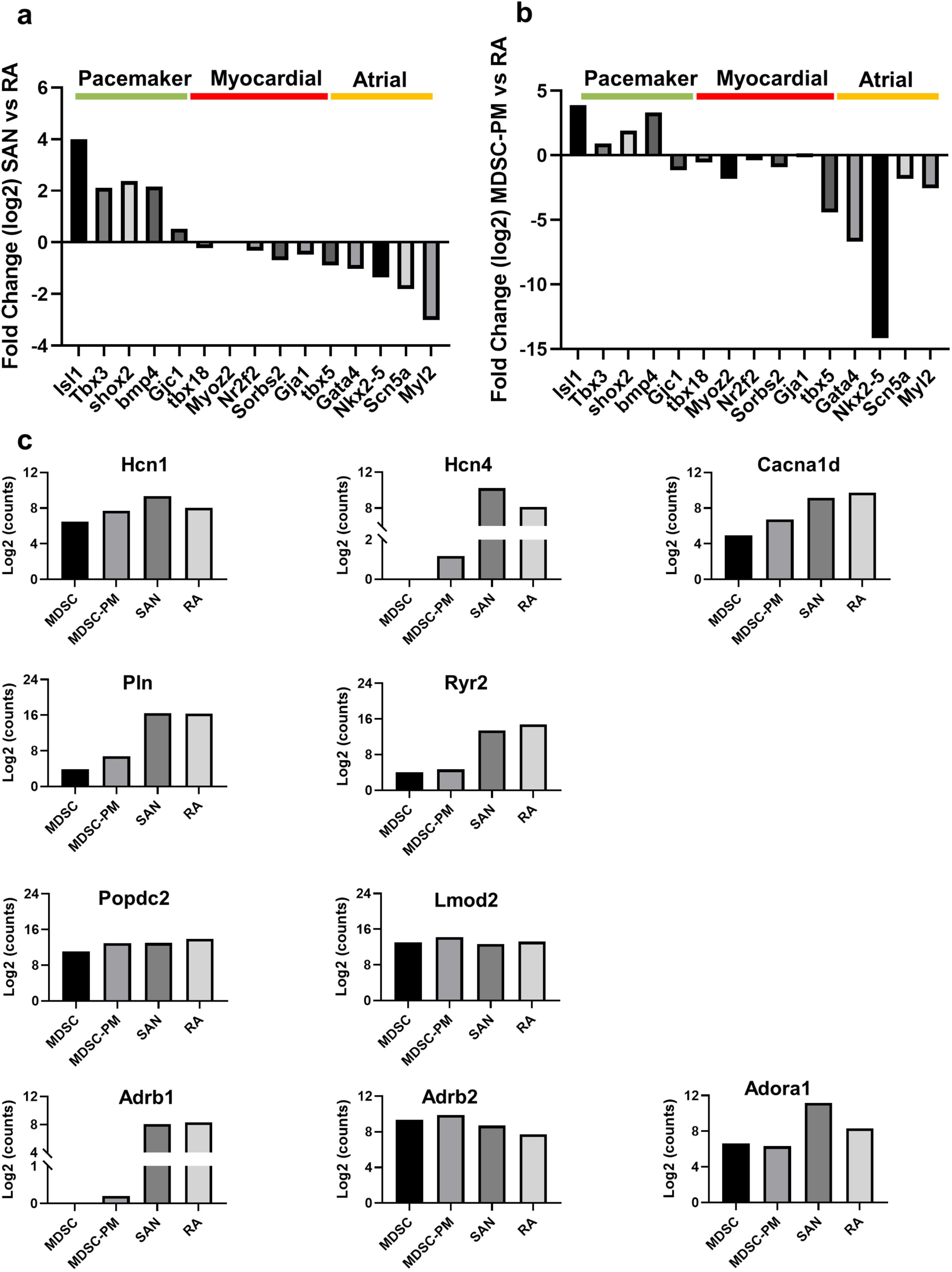
Comparative analysis of RNAseq-profiled gene expression in MDSC-PM and native SAN and RA tissue. (**a-b**) Differential expression of critical marker genes involved in the specification of SAN tissue (pacemaker), working myocardium (myocardial) and right atrium (atrial). In panel (a) and (b), differential expression is expressed as the Log_2_ of the ratio of absolute pseudo-counts-RLE of a given gene between the SAN and the right atrium (a), or between MSDC-PM and the right atrium (b). Positive values indicate increased expression, while negative indicate reduced expression. (**c**) Expression of genes coding for key ion channels, membrane receptors and regulators involved in pacemaker activity in undifferentiated MDSC, differentiated MDSC-PM, the SAN and the right atrium (RA).

Another hallmark of cardiac pacemaker cells is their expression of the channel HCN4 ^28^, which is responsible for generating the “funny” current (*I*_*f*_) ^29^. RT-PCR analysis of MDSC-PMs (Fig. 2b, Supplementary Table 1) showed that they express the pore-forming subunits of HCN4 and Ca_v_1.3 channels, both of which are required for normal cardiac pacemaker activity in mice ^30-34^ and humans ^35,36^. MDSC-PMs also express the pacemaker-specific gap junction protein Cx45 ^14,37^ and the cardiac contractile proteins α-actinin and Troponin I (Fig. 2b). The co-expression of these proteins in the same pacemaker-like MDSC-PM was examined by double-immunostaining of MDSC-PMs differentiated from MDSCs isolated from wild type mouse muscle, In order to show differential and specific staining for the protein markers analyzed, three microscopic fields with well identifiable pacemaker-like myocytes are shown with co-expression of HCN4 with cardiac troponinI (Fig.2c left panels), Ca_v_1.3 with α-actinin (Fig.2c middle panels) and Cx45 with α-actinin (Fig.2c right panels). Given that HCN proteins, Ca_v_1.3, and Cx45 are also expressed in neural cells, cells co-expressing cytoskeletal markers of striated muscle cells (cardiac Troponin I or sarcomeric α-actinin) with these markers demonstrate a cardiac pacemaker phenotype (Fig.2c). Of note, in the panels showing co-staining for α-actinin and Cx45, a typical neural cell on the right expressed Cx45 but not α-actinin. As further evidence, Fig. 2d shows cells co-stained for Ca_v_1.3 and Islet1, a marker only expressed in pacemaker cells in adult heart^27^, and includes a phase-contrast image of the MDSC-PM undergoing sustained beating and filmed before fixation (see corresponding Supplementary video 4). Of note, the beating pacemaker-like myocytes differentiated from MDSCs expressed none of the 4 myogenic transcription factors, MyoD, Myf5, myogenin or Mrf4 as probed by in situ immunofluorescence and confirmed by positive staining on differentiated primary myoblasts and myotubes (data not shown).

To further examine the expression pattern of genes linked to pacemaker activity, the expression levels of key markers in MDSC-PMs differentiated *in vitro* was compared to expression of the same markers in native sinoatrial (SAN)^38^ and right atrial (RA) tissues. Fig. 3 illustrates the relative expression levels of 25 critical mRNAs encoding transcription factors, ion channels, connexins and cardiac cytoskeletal proteins in native cardiac cells and in MDSC differentiated *in vitro* into MDSC-PM. Consistent with previous reports^38,39^, the native SAN tissue expressed high levels of transcription factors involved in the differentiation of pacemaker tissue, while expression of myocardial and atrial markers was low (Fig. 3a). Conversely, the expression of pacemaker markers was shared between MDSC-PM and native SAN (Fig. 3b) whereas typical markers of atrial myocytes, including the voltage-gated Na^+^ channel Scn5a (Na_v_1.5) and Myl2 where down-regulated in MDSC-PM similarly to native SAN in comparison to native atrial tissue (Fig. 3b)Analysis of f-(HCN) channel expression reveals an increase in the expression of mRNAs coding for HCN1 and HCN4 (Fig. 3c) upon differentiation of MDSC in MDSC-PM. In contrast there was no evidence for expression of atrial myocyte markers in MDSC-PM (Fig. 3b). These data show that MDSC differentiated into beating pacemaker-like cells do not induce expression of ventricular- or atrial-specific differentiation markers.

Importantly, *in vitro* differentiation of muscle-derived stem cells into MDSC-PMs was not a peculiarity of mouse muscle stem cells. It was also observed in muscle stem cells isolated from adult Sprague Dawley rat muscle (Supplementary Fig. 2 and Supplementary video 5), and both the differentiation kinetics and co-expression of pacemaker markers including HCN4 with α-Actinin (shown in Supplementary Fig. 2) were similar to that in mice.

We next tested MDSC-PMs for the functional properties of native SAN pacemaker myocytes using current-clamp recordings (Fig. 4). Patch-clamp recordings from isolated beating MDSC-PMs myocytes revealed spontaneous repetitive action potentials (Fig. 4a) at rates comparable to those produced by native SAN myocytes^33^, in spite of a significantly higher slope of the diastolic depolarization in native pacemaker myocytes, probably because of a shorter action potential duration in MDSC-PM (Supplementary Fig. 3a-c *and* Fig. 3i-l). The membrane capacitance of differentiated MDSC-PMs was lower than that of atrial myocytes, but was similar to that of SAN myocytes (Supplementary Fig. 3d). We then compared the parameters of action potentials recorded in MDSC-PM to those of native SAN and atrial myocytes (Supplementary Fig. 3). The maximum diastolic potential, action potential threshold and upstroke velocity were similar between recordings of MDSC-PMs and native SAN myocytes (Supplementary Fig. 3e-g). However, the action potential of MDSC-PMs showed significantly smaller amplitude and shorter duration than that of native atrial and SAN myocytes (Supplementary Fig. 3g *and* h). Whether contracting myocytes generate automatic spontaneous intracellular Ca^2+^ ([Ca^2+^]_i_) transients similar to those produced by native mouse SAN pacemaker myocytes was tested by loading MDSC-PMs with Fluo-4-AM. Line-scan images revealed spontaneous [Ca^2+^]_i_ transients (Fig. 4b). [Ca^2+^]_i_ transients recorded in MDSC-PM showed significantly lower amplitude and longer duration than those of native SAN myocytes (Supplementary Fig. 4). Images were then recorded before and after the cultures were perfused with epinephrine or acetylcholine, to determine whether the frequency of spontaneous [Ca^2+^]i transients was regulated in opposite directions by the activation of β-adrenergic ^34^ and muscarinic ^40^ receptors. Indeed, epinephrine (2 nM) accelerated, whereas acetylcholine (50 nM) reduced, the frequency of spontaneous [Ca^2+^]_i_ transients. Ryanodine receptors (RyRs) are also involved in SAN pacemaker activity (reviewed in ref. ^41^). Perfusion of MDSC-PMs with ryanodine (Fig. 4b) reduced the amplitude and prolonged the duration of spontaneous [Ca^2+^]i transients (Supplementary Fig. 5). In addition, ryanodine perfusion led to a near-complete arrest of spontaneous [Ca^2+^]_i_ transients (Fig. 4b). Thus, RyRs appear to play a major role in the automaticity of MDSC-PMs, similar to that in native SAN pacemaker myocytes ^41^.

**Fig. 4.**
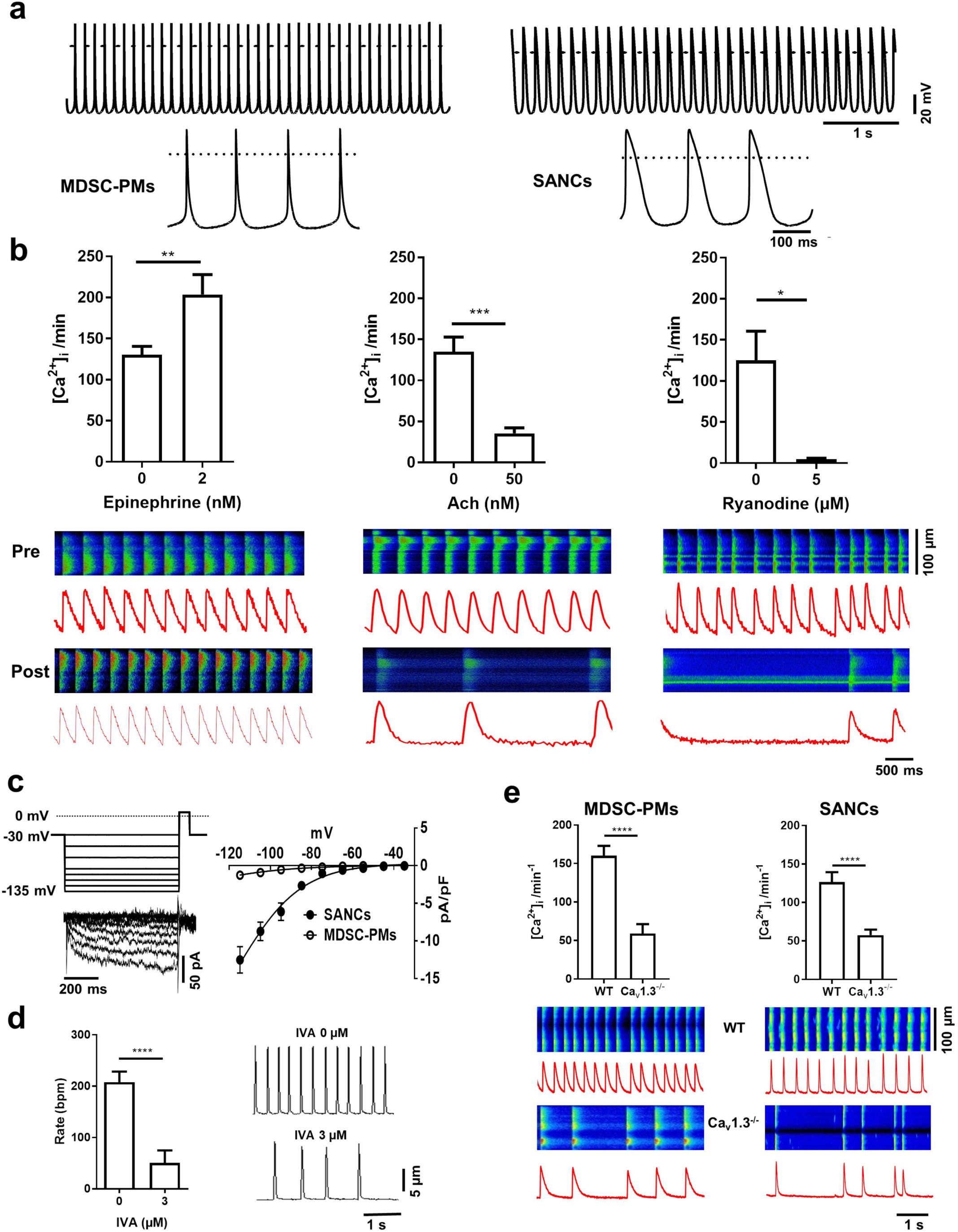
Electrophysiology and Ca^2+^ signaling in pacemaker-like MDSC-PMs differentiated *in vitro*. (**a**) Representative traces of action potentials recorded from *in vitro*-differentiated beating MDSC-PMs (*left panel*) and native SAN pacemaker myocytes (*right panel*). Insets show action potentials at expanded scale. (**b**) Chronotropic response of Fluo-4-loaded MDSC-PMs, measured as the change in frequency of spontaneous [Ca^2+^]_i_ transients pre-(*top line*) and post-(*bottom line*) perfusion with the following agonists: epinephrine, 2 nM (*left panel*: n=12); acetylcholine (Ach) 50 nM (*middle panel*: n=11); and ryanodine, 5 µM (*right panel*: n=5). Corresponding representative confocal line-scan images of [Ca^2+^]_i_ transients in MDSC-PMs. Light intensity integrals (red traces) are shown underneath each scan. Statistical significance was tested using paired t-test. *p<0.05, **p<0.01. Error bars indicate s.e.m.. (**c**) Voltage protocol, sample current traces *(left panel*) and averaged current-to-voltage relationship of *I*_*f*_ (*right panel*) recorded in differentiated MDSC-PMs (peak *I*_*f*_ density at –135 mV: 2.5 ± 0.2 pA/pF, n=11) and native SAN myocytes (23±3 pA/pF, n=22). *I*_*f*_ was evoked by applying hyperpolarizing voltage steps from a holding potential of –30 mV.(**d**) Averaged spontaneous contraction rates (*right panel*) and corresponding edge-detection recording samples (*left panel*) of *in vitro*-differentiated MDSC-PMs under control conditions, without ivabradine (0 IVA) and during perfusion with ivabradine (IVA 3 μM, n=15). Statistical significance was tested using paired t-test. ****p<0.0001. Error bars indicate s.e.m.. (**e**) Spontaneous [Ca^2+^] transients in MDSC-PMs (*left panel*) derived from wild-type (WT, *top line*) and Ca_v_1.3^-/-^ (*bottom line*) mice, compared to spontaneous [Ca^2+^]_i_ transients measured in isolated native SAN cells (SANCs, *right panel*) from wild-type (*top line*) and Ca_v_1.3^-/-^ (*bottom line*) mice. Light intensity integrals (red traces) are shown underneath each scan. Statistical significance was tested using the unpaired t-test. ****p<0.0001. Error bars indicate s.e.m..

**Fig. 5.**
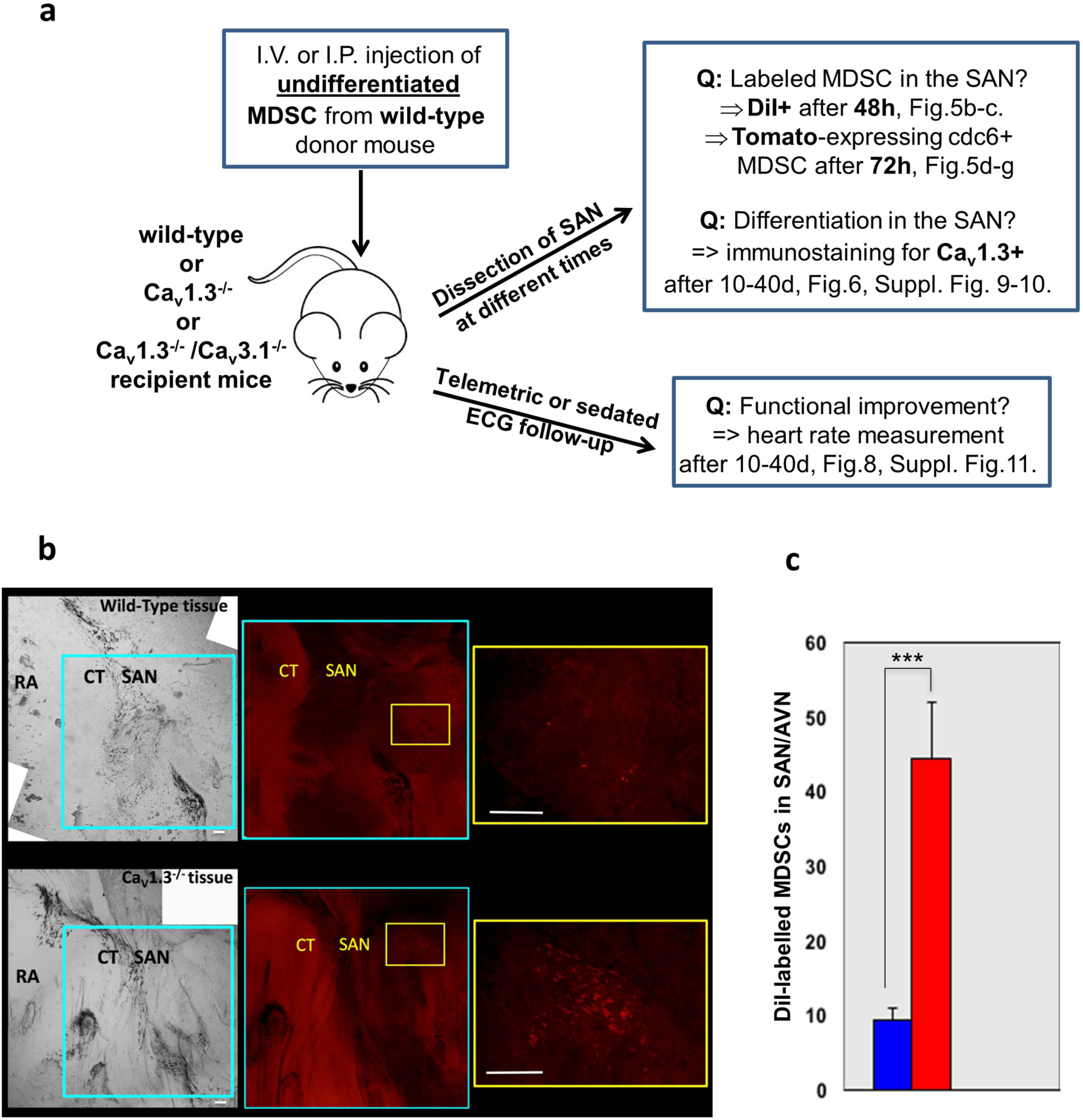
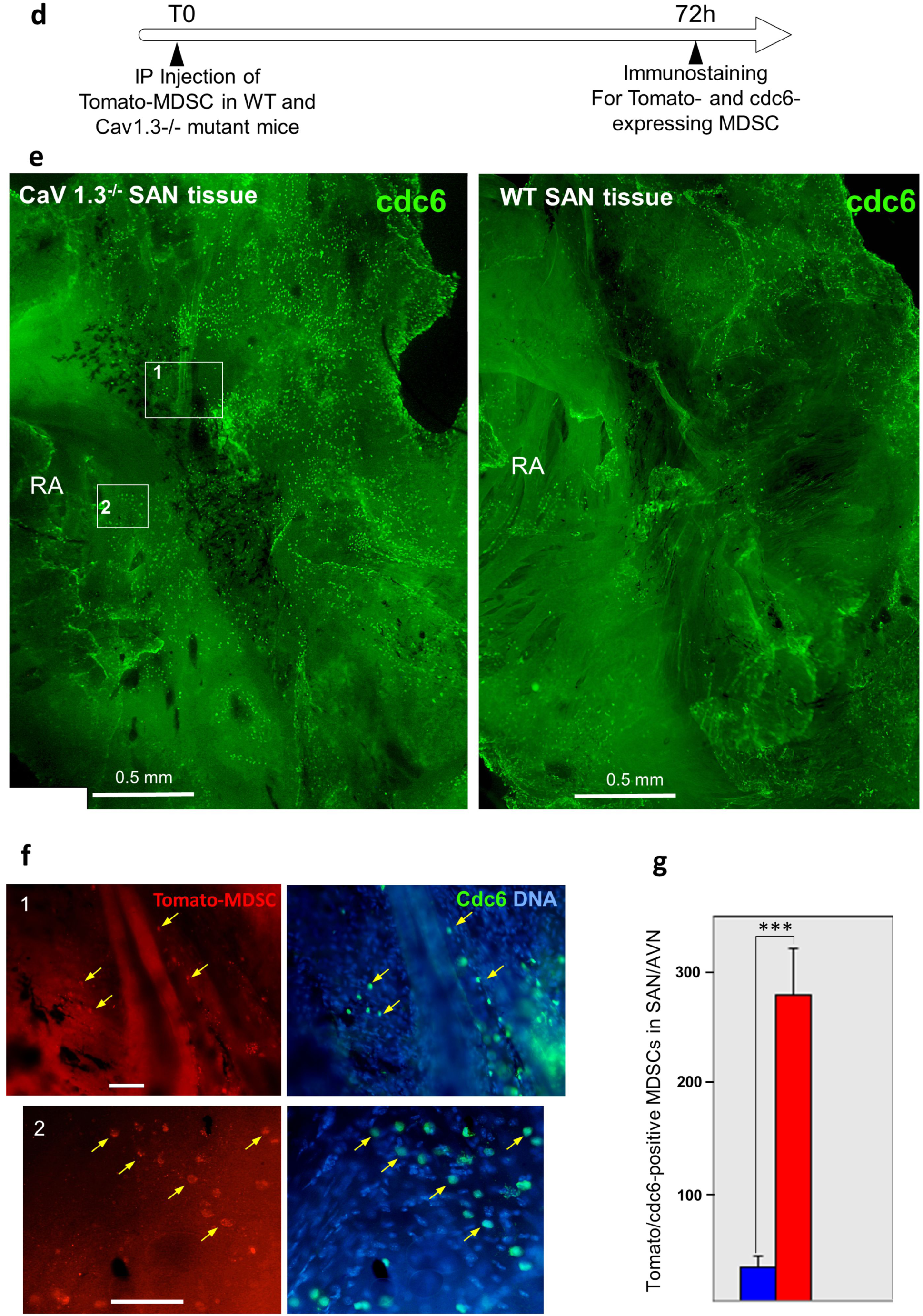
Summary of *in vivo* experiments with bradycardic mutant mice and engraftment to the SAN by MDSCs. (**a**) Schematic representation of assays testing the early homing and reparative potential of MDSCs *in vivo*. (**b**) Assays of MDSC migration and early engraftment to SAN tissue, in age- and sex-matched wild-type and Ca_v_1.3^-/-^ mice 48 hours after I.P. injection with 3×10^5^ MDSCs pre-labelled with the membrane marker DiI for cell tracking. Top panels show SAN tissue from a male wild-type mouse injected with DiI-MDSCs. Bottom panels show SAN tissue from a male Ca_v_1.3^-/-^ mouse injected with DiI-MDSCs. Shown are representative bright-field images at 60x and close-ups (areas shown by frames) of red fluorescence (DiI). RA: right atrium; CT: crista terminalis; SAN: sino-atrial node area. Bar = 200 µm. (**c**) Histogram representation of the quantification of engrafted DiI-labelled MDSCs in SAN tissue from 3 wild-type (blue) and 3 Ca_v_1.3^-/-^ (red) mice Statistical significance was tested using the unpaired t-test. ***p<0.001. Error bars indicate s.e.m.. (**d**-**g**) Increased engraftment after 3 days (72 hr) by cdc6-positive MDSCs to the SAN of mutant Ca_v_1.3^-/-^ mice. (**d**) Schematic representation of MDSC transplantation in wild-type (WT) and Ca_v_1.3^-/-^ mice and immunostaining of SAN tissue after 3 days to visualize MDSCs engraftment. (**e**) Immunostaining for cdc6 (cell division cycle 6 involved in the control of early steps of DNA replication) in wild-type (WT, right panel) and Ca_v_1.3^-/-^ (left panel) SAN tissue showing large field distribution of the signal with the position of 2 framed fields shown at higher magnification in the bottom panels. RA: Right atrium. bar: 500 µm. (**f**) Enlarged panels 1 and 2 showing co-staining for tomato-expressing MDSCs (red, left panels) and cdc6-positive cells with Hoechst DNA staining (green and blue, right panels), arrowed are some of the cells co-expressing Tomato and cdc6. Bars: 50 µm. (**g**) Histogram representation of engrafted tomato-expressing MDSCs counted in the SAN area after 3 days in 3 independent transplantation experiments in wild type-mice (blue bar) and in Ca_v_1.3^-/-^ mice (red bar). Statistical significance was tested using the unpaired t-test. ***p<0.001. Error bars indicate s.e.m..

Consistent with HCN4-positive staining (Fig. 2), MDSC-PM myocytes displayed the *I*_*f*_ current (Fig. 4c), which is involved in the pacemaker activity of native SAN myocytes ^29,42^. The density of *I*_*f*_ in MDSC-PM was lower than that recorded in native SAN myocytes (Fig. 4c). We then recorded the spontaneous contraction frequency of beating cells by using a cell edge detection device (see methods). The selective *I*_*f*_ inhibitor ivabradine (IVA) ^43^ reduced the frequency of spontaneous cell contraction by more than 75%, demonstrating that, despite its lower density, the *I*_*f*_ is a key determinant of the spontaneous activity of MDSC-PMs (Fig. 4d).

We then employed confocal imaging of [Ca^2+^]i release to investigate further the mechanism underlying pacemaking of MDSC-PMs. We previously showed that loss of Ca_v_1.3 channels slows automaticity and leads to arrhythmia in native SAN myocytes ^30,34^. To test whether MDSC-PMs differentiated from Ca_v_1.3 knockout (Ca_v_1.3^-/-^) mice reproduced the phenotype of native Ca_v_1.3^-/-^ SAN myocytes, we compared their beating frequency to that of MDSC-PMs derived from wild-type MDSCs. As shown Fig. 4e, the rate of spontaneous [Ca^2+^]_i_ transients in MDSC-PMs differentiated from Ca_v_1.3^-/-^ MDSCs was significantly slower and their frequency less regular than those in MDSC-PMs from wild type MDSCs, consistent with findings from our previous work on native SAN myocytes from Ca_v_1.3^-/-^ mice ^34,40^.

Together, these data show that MDSCs are capable of differentiating *in vitro* into beating pacemaker-like myocytes with all the specific markers and electrophysiological properties of native cardiac pacemaker cells. Furthermore, *in vitro* differentiation of MDSCs derived from the muscle of bradycardic Ca_v_1.3^-/-^ mutant mice with SAN dysfunction reproduced in MDSC-PMs defects of cellular automaticity similar to native knockout cells. These results also demonstrate that Ca_v_1.3 channels play a key role in the pacemaker activity of MDSC-PMs, consistent with *in vivo* data from Ca_v_1.3^-/-^ mice ^40,44,45^ and from humans with a congenital loss-of-function mutation in Ca_v_1.3 channels ^36,46^.

### *In vivo* analysis: MDSCs injected into mice lacking Ca_v_1.3 and Ca_v_3.1 channels migrate and home to the SAN where they differentiate resulting in improved heart rate

To test the ability of MDSCs to differentiate *in vivo* into pacemaker cells and their potential therapeutic effects, we employed the Ca_v_1.3^-/-^ deficient mouse, a well-characterized model of human SAN dysfunction ^36,40^, as recipients in stem cell transplantation experiments. Both embryonic stem cells (ESCs) and induced pluripotent stem cells (iPSC) can be differentiated into SAN-like cardiomyocytes *in vitro* and have been shown to provide promising short-term “biological” pacemaker activity in the ventricle ^47-50^. However, the high proliferation rate and associated tumorigenicity of these cell types preclude their direct use without prior differentiation for *in vivo* cell therapy. Mouse ESCs grafted subcutaneously onto 8-week-old immunodeficient SCID mice (n=2) induced tumour outgrowth at the site of the graft and required euthanasia after 3 weeks. In contrast, we found no evidence for a long-term teratogenic or tumorigenic potential of MDSCs. No tumours formed in mice (n=4) engrafted with MDSCs, and the animals remained tumour-free for over 4 months after either sub-cutaneous (Supplementary Fig. 6a) or intraperitoneal (I.P.) transplantation (data not shown, n=6). In addition, RT-PCR analysis (n=4) revealed that MDSCs express 3 of 4 pluripotency transcription factors (Sox2, Nanog, Lin28) but not the major one, Pou5F1/Oct4 (Supplementary Fig. 6b) a result confirmed by RNAseq analysis on triplicate MDSC samples (not shown). The lack of teratoma formation after MDSC injection supports the direct use of these stem cells *in vivo*, without need to induce them into differentiation.

**Fig. 6.**
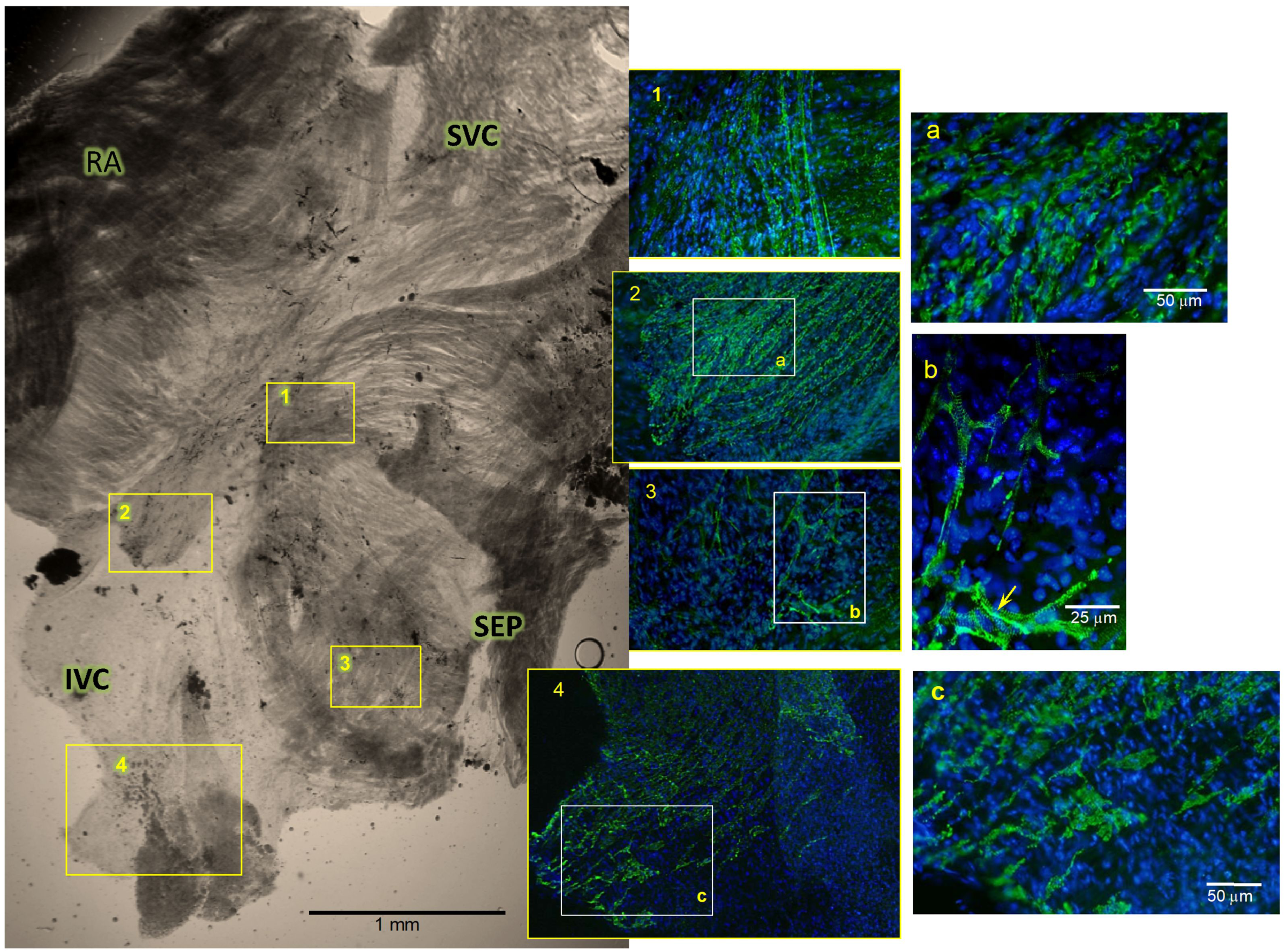
Engraftment and differentiation of wild-type MDSCs in the SAN tissue of Ca_v_1.3^-/-^ mutant mice. SAN tissue of Ca_v_1.3^-/-^ mutant mice injected (I.V.) with MDSCs, shown at 40 days post-injection. Immunofluorescence labelling for Ca_v_1.3 (green) was carried out on whole-mount SAN tissue together with Hoechst DNA staining as described in Methods. Shown are bright-field images of the whole SAN region, with the right atrium (RA, cut before final mounting), superior and inferior vena cava (SVC and IVC) and inter-atrial septum (SEP) marked. Higher-magnification immunofluorescence images of areas marked 1-4 in the bright-field image show arrays of individual Ca_v_1.3-positive cells with typical patchy membrane staining. Further magnification of areas **a, b**, and **c** show membrane-associated and striated patterns, with enhanced staining at cell junctions (arrow in **b**). Similar results were obtained from 5 independent MDSC injection experiments into Ca_v_1.3^-/-^ mice.

Tracking DiI membrane-labelled MDSCs 24 and 72 hours after I.P. or I.V. injection, indicates a similar distribution to major filtration organs (e.g., liver and kidney) and lymph organs (spleen and lymph nodes) in wild-type and mutant recipient mice (Davaze, Mitutsova et al. manuscript in preparation). Similar analyses of I.P.-injected, labelled-MDSCs showed enhanced MDSC homing to organs with targeted injury, such as streptozotocin-damaged pancreatic islets ^51^. Thus, MDSCs used directly as stem cells *in vivo* without prior differentiation have the ability to migrate and home to a variety of organs. This property was exploited to test the capacity of MDSCs to migrate to the SAN tissue, differentiate *in vivo*, and restore compromised heart automaticity in our mouse models with SAN dysfunction, as represented in Fig.5a. To first assay the early homing of MDSC to wild-type and mutant SAN tissue, MDSCs labelled with the membrane-binding cell tracker DiI-CM were transplanted, by I.V. or I.P. injection, into Ca_v_1.3^-/-^ mutant mice carrying sinus node dysfunction associated with atrioventricular conduction blocks, as well as into age-matched wild-type counterparts. At 48 hours after injection, a significant number of labelled stem cells were found implanted in the SAN tissue of recipient Ca_v_1.3^-/-^ mice (Fig.5b), whereas only few were present in the SAN of wild-type mice (Fig.5b *and* c); this finding was reproduced in 4 wild-type and 4 Ca_v_1.3^-/-^ mutant mice using the same MDSC preparation. Indeed, as counted from these 4 transplantation experiments, the number of DiI-labelled MDSC in Ca_v_1.3^-/-^ SAN was 5-fold higher than in wild-type counterparts (Fig. 5c). To further probe this increased engraftment to the mutant SAN, we injected MDSC obtained from mice globally expressing membrane-bound Tomato fluorescent marker^52^ and tested their engraftment 3 days after I.P. transplantation (Fig. 5d-g). Dissected SAN tissues were co-stained for Tomato and cdc6, cell cycle regulator of DNA synthesis^53^, to reveal MDSC that are competent to divide. Staining for cdc6 shown in Fig.5e shows large amounts of positive cells in the SAN area of Ca_v_1.3^-/-^ hearts compared to wild-type SAN tissue. Co-staining for Tomato-expressing MDSC (Fig.5f) shows that most of the cdc6-expressing cells also expressed Tomato, corresponding to replication-competent MDSCs. Quantification of co-stained Tomato and cdc6-expressing cells from 3 different experiments showed 10-fold higher cell number engrafted to Ca_v_1.3^-/-^ SAN, than to wild-type counterparts (Fig.5g). These results show that I.P.-injected MDSC effectively migrate to the SAN tissue where they display increased homing to the Ca_v_1.3^-/-^ mutant SAN compared to that in wild type SAN tissue. In addition, the extensive expression of cdc6 in engrafted MDSCs shown 3 days after injection is indicative of their sustained capacity to divide after engraftment. Such increased homing of MDSCs in mutant SAN tissue may be related to the SDF-1–CXCR4 chemotactic axis, a major pathway in the regulation of stem-cell migration and homing ^54,55^. Indeed, comparison of the expression of SDF1 and CXCR4 levels in SAN tissue isolated from the two groups of mice showed that SAN tissue from Ca_v_1.3^-/-^ mice expressed higher levels of both SDF1 and CXCR4 than did SAN tissue from wild-type counterparts. The RT-PCR with quantitative analysis (Supplementary Fig. 7a-b) shows a 2- and 3-fold increased expression of CXCR4 and SDF1 respectively in Ca_v_1.3^-/-^ SAN tissue. This data was further confirmed at the protein level by Western blot analysis of the SAN tissue from wild type, single Ca_v_1.3^-/-^ and double Ca_v_1.3^-/-^ / Ca_v_3.1^-/-^ mutant. In both mutants the SAN displayed significantly higher levels of CXCR4 and SDF1 than the tissue from wild type mice (shown in Supplementary Fig. 7c) with ca. 2.5 fold and 4 fold increases in CXCR4 and SDF1 levels respectively in mutant SAN tissue as quantified relatively to GAPDH levels in the histogram represented in Supplementary Fig.7d.

**Fig. 7.**
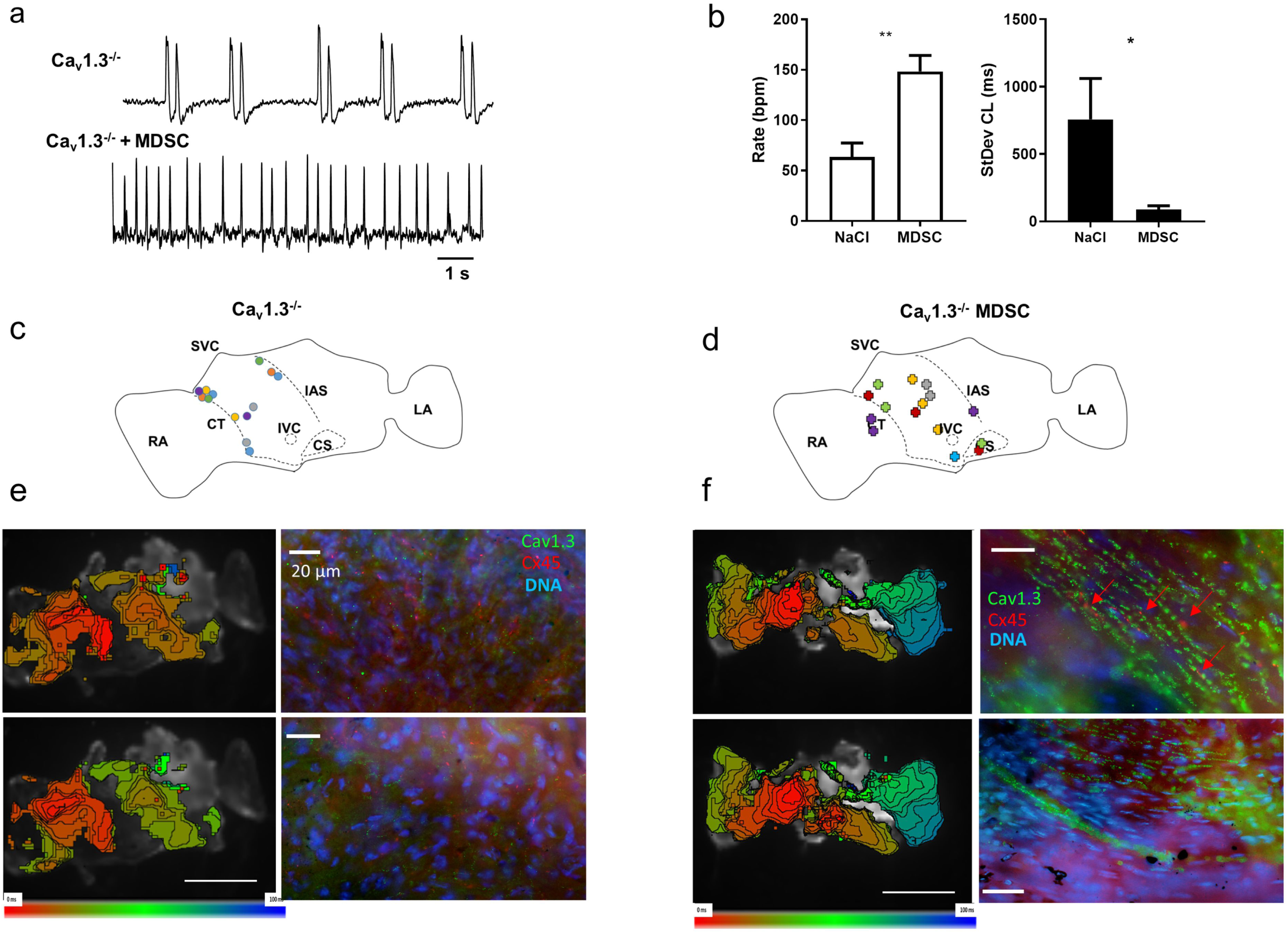
Optical mapping of pacemaker impulse in the sinus node of MDSC injected Ca_v_1.3^-/-^ mice. (**a**). Sample traces of optically recorded spontaneous action potentials in Ca_v_1.3^-/-^ sinus nodes from mice injected with vehicle (NaCl, upper panel, n=6), or MDSCs (bottom panel, n=6). (**b**). Corresponding averaged rates of spontaneous action potentials (left) and the standard deviation (variability) of the inter beat duration (right) in atrio-sinus preparations injected with vehicle (black bars), or MDSCs (gray bars). Statistical significance was tested using the unpaired t-test. *p<0.05, **p<0.01. Error bars indicate s.e.m.. Position of the pacemaker leading sites in Ca_v_1.3^-/-^ sinus nodes injected with vehicle (**c**), or MDSC (**d**). Each color corresponds to an individual atrio-sinus preparation. Multiple points of a given color indicate co-existence of different leading sites in a given preparation. (**e**-**f**). Samples of activation maps derived from optical mapping recordings of atrio-sinus preparations from vehicle-injected (**e**) and MDSC injected (**f**) Ca_v_1.3^-/-^ mice. The right panels show corresponding close-up views of pacemaker leading sites co-stained with anti-Ca_v_1.3 and anti-Cx45 primary antibodies. Arrows indicate contact points between cells. Nuclei are stained in blue. Statistics: Abbreviations: RA, right atrium; CT, crista terminalis; SVC, superior vena cava; IVC, inferior vena cava; CS, coronary sinus; IAS, inter-atrial septum; LA, left atrium. Statistics: unpaired t-type test.

We determined whether SAN engraftment of injected wild type MDSC was followed by their differentiation into MDSC-PM. To this aim, we isolated intact SAN tissue from the hearts of control uninjected wild type, NaCl injected (mock-transplanted) Ca_v_1.3^-/-^ and Ca_v_1.3^-/-^ mice 30 to 40 days after MDSC injection, to compare the expression of the Ca_v_1.3 pore-forming α1 subunit using whole-mount immunostaining for cells expressing Ca_v_1.3. Indeed, we reasoned that since Ca_v_1.3 α1 subunit is absent in the SAN tissue originating from recipient Ca_v_1.3^-/-^ SAN^44,56^, *in situ* differentiation of MDSC-PM would have been revealed by the presence of Ca_v_1.3 positive cells in the Ca_v_1.3 knockout SAN. The SAN of wild type mice displayed continuous widespread Ca_v_1.3 staining in the entire SAN region (Supplementary Fig. 8). Staining for Ca_v_1.3 was specific because it was absent in the SAN of control mock-transplanted Ca_v_1.3^-/-^ mice (n=5, Supplementary Fig. 9). As shown in Fig. 6 and in Supplementary Fig. 9, the SAN of MDSC transplanted Ca_v_1.3^-/-^ mice was also positive for Ca_v_1.3 staining. In comparison to the wild type SAN, Ca_v_1.3 staining of recipient SANs was characterized by the presence of arrays of positive cells (zoomed panels Fig. 6a and 6c with 49 and 54 Ca_v_1.3 -expressing cells, respectively) showing membrane and striated patterns of staining for Ca_v_1.3 similar to those for native mouse SAN pacemaker cells ^56^, including increased staining at junctions between cells with typical pacemaker morphology (arrow in Fig. 6b). The effectiveness of the differentiation of MDSCs into Ca_v_1.3-expressing myocytes and their long-term colonisation was further confirmed by clear detection of Ca_v_1.3-stained cells in the SAN of Ca_v_1.3^-/-^ /Ca_v_3.1^-/-^ double-mutant mice 5 months after I.V. injection of wild-type MDSCs (Supplementary Fig. 10).

**Fig. 8.**
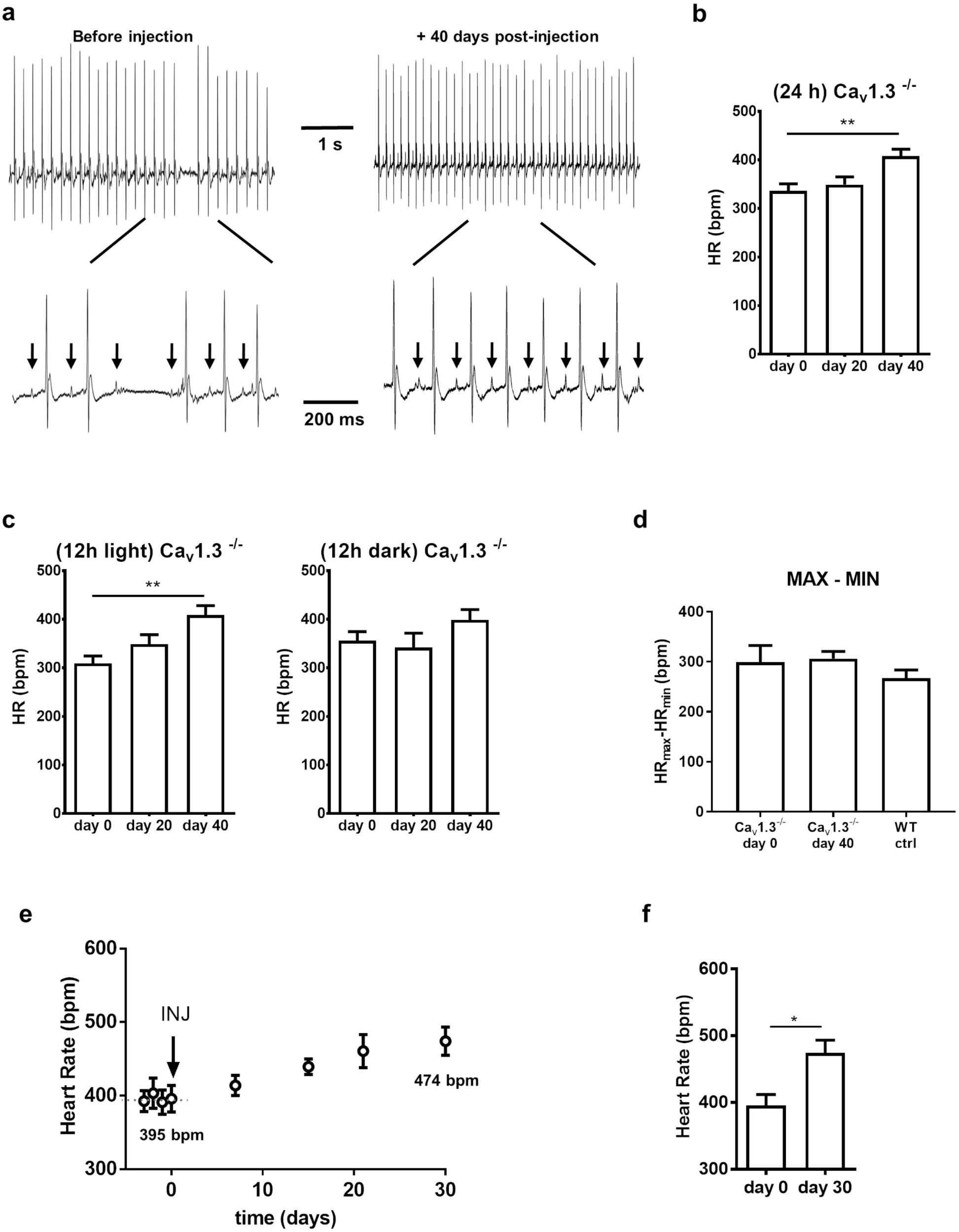
*In vivo* recordings of heart rate in Ca_v_1.3^-/-^ single-mutant and Ca_v_1.3^-/-^/Ca_v_3.1^-/-^ double-mutant mice before and after transplantation of MDSCs from wild-type muscle. (**a**) Samples of *in vivo* telemetric ECG recordings from a freely moving Ca_v_1.3^-/-^ mouse before and after systemic transplantation (I.V. injection) of MDSCs. (**b**) Quantitation of 24-hour telemetric recordings of heart rate in n=6 Ca_v_1.3^-/-^ mice, before and up to 40 days after systemic transplantation of MDSCs. (**c**) Quantitation of 12-hour recordings during light (left panel) and dark (right panel) period of heart rate (see Table 1) in n=6 Ca_v_1.3^-/-^ mice, before and up to 40 days after systemic transplantation of MDSCs. (**d**) Relative degree of heart rate regulation expressed as the difference between the highest (MAX) and lowest (MIN) heart rate. Statistical significance was tested using analysis of variance (ANOVA) followed by Kruskal-Wallis multiple comparisons test. *p<0.05, **p<0.01. Error bars indicate s.e.m.. (**e**) Quantitation of *in vivo* telemetric heart rate recordings in Ca_v_1.3^-/-^/Ca_v_3.1^-/-^ double mutant mice before and after systemic transplantation of MDSCs (I.P. injection). (**f**) Quantitation of heart rate recorded in Ca_v_1.3^-/-^/Ca_v_3.1^-/-^ mice before (day 0) and on day 30 after I.P. transplantation of MDSCs from wild-type mice. Statistical significance was tested using the paired t-test. *p<0.05. Error bars indicate s.e.m

We then investigated whether engraftment of MDSC resulted in electrical integration of differentiated MDSC-PM to affect SAN automaticity of Ca_v_1.3^-/-^ mice. To this end, we employed optical mapping of membrane voltage to record the pacemaker impulse and the site of origin of automaticity in isolated SAN/atria preparations from Ca_v_1.3^-/-^ mice 40 days after injection of vehicle (NaCl) or MDSC (Fig. 7). Optical mapping of mock-transplanted Ca_v_1.3^-/-^ SANs showed low-rate automaticity and high inter beat variability (Fig. 7a *and* b). In contrast, automaticity of MDSC-transplanted Ca_v_1.3^-/-^ SANs was characterized by ∼2-fold higher pacemaking rate and reduced inter beat variability (Fig. 7a *and* b). We thus hypothesized that electrical integration of MDSC-PM generated new leading pacemaker sites expressing Ca_v_1.3 that improves pacemaker activity in the SAN of transplanted Ca_v_1.3^-/-^ mice. We thus correlated the position of the leading pacemaker sites to the expression of Ca_v_1.3 and Cx45 in mock- and MDSC-transplanted Ca_v_1.3^-/-^ SANs (Fig. 7c-f). Staining of leading pacemaker sites in mock-transplanted Ca_v_1.3^-/-^ SANs showed Cx45 positive cells, but no detectable Ca_v_1.3 staining (Fig. 7e). In contrast, imaging of the SAN leading pacemaker sites in MDSC transplanted Ca_v_1.3^-/-^ mice showed cells co-expressing Ca_v_1.3 and Cx45 (Fig. 7f). Particularly, Cx45 staining was observed along cell borders in contact points between cells expressing Ca_v_1.3 and Ca_v_1.3 negative cells, suggesting electrical coupling between donor-derived MDSC-PM and pacemaker myocytes of the recipient SAN.

Whether colonization of the SAN by MDSCs and their differentiation in MDSC-PM in bradycardic mutant mice was accompanied by an improvement in heart rhythm *in vivo* was determined by analysing heart rate during 24-hour periods. To this end, telemetric ECG recordings were made in freely moving Ca_v_1.3^-/-^ mice, before and at different times after the injection of undifferentiated wild-type MDSCs (Fig. 8a-b). Before injection, the heart rhythm of Ca_v_1.3^-/-^ mice was characterized by SAN bradycardia (low PP interval) and frequent occurrence of SAN pauses indicative of underlying SND in accordance with our previous study^40^. Ca_v_1.3^-/-^ mice experienced also 2^nd^ degree atrioventricular blocks (Fig. 8a and Table 1)^40^. By 40 days after systemic transplantation of muscle-derived stem cells, the MDSC-injected mice had a significantly higher (∼25%, Fig. 8b) heart rate than before the procedure (Fig. 8b), indicating that the SAN pacemaker activity had improved. Analysis of heart rate during the 12-hour day time and night time indicated that the relative effect of MDSC injection was maximal during resting day time (∼32%, Fig. 8c, Table 1). Heart rate elevation in Ca_v_1.3^-/-^ mice was due to improvement of SAN rate (PP interval). No significant differences were observed in heart rate and the other ECG parameters in wild-type mice (Table 1). Similarly, no significant changes in heart rate were observed following implantation of only a ECG transmitter (no MDSCs; Supplementary Fig. 11a) or when control wild-type mice were injected with MDSCs (Table 1). In addition, the difference between maximal and minimal heart rates were similar in wild-type and Ca_v_1.3^-/-^ mice injected with MDSC (Fig. 8d), indicating that transplantation of MDSCs did not affect the relative degree of heart rate regulation. All control animals were kept under identical conditions as MDSC-injected Ca_v_1.3^-/-^ mice. Measurement of QT, rate-corrected QT (QTc) and QRS ECG intervals showed that MDSC injection did not affect ventricular repolarization, and no episodes of ventricular arrhythmia or sudden death were observed in MDSC-injected Ca_v_1.3^-/-^ and wild-type mice (Table 1). We then tested whether injection of MDSC could improve the heart rate in mice lacking both Ca_v_1.3 and Ca_v_3.1 channels (Ca_v_1.3^-/-^ /Ca_v_3.1^-/-^)^57^. This mouse model presents with strong SND hallmarks and highly variable SAN rate. If MDSC injection ameliorated the heart rate of Ca_v_1.3^-/-^/Ca_v_3.1^-/-^ mice, this would have been reflected as an increase in SAN rate and in a reduction in the inter beat variability. Indeed, upon injection of wild-type MDSCs, the heart rates of Ca_v_1.3^-/-^/Ca_v_3.1^-/-^ mice were clearly improved within 30 days (as measured during the active night-time period: +20%) (Fig. 8e-f, Supplementary Table 2). Heart rate of double-mutant Ca_v_1.3^-/-^/Ca_v_3.1^-/-^ and wild-type mice was also recorded under anesthesia 10 to 20 days after injection of MDSCs and in mice monitored over 5 months after 3 repeated injections of MDSCs (see example of long term engraftment in Supplementary Fig. 10). This recording of sedated mice revealed a significant increase in heart rate in the double-mutant mice by 10 days (+28%, Supplementary Fig. 11b) and a decrease in the inter beat standard deviation after 20 days (Supplementary Fig. 11c), whereas similar injection of MDSCs into wild-type mice did not result in significant changes in either heart rate or inter beat standard deviation (Supplementary Fig. 11b-c). The number of SAN pauses and AV blocks after MDSC injection into either single Ca_v_1.3^-/-^ or double Ca_v_1.3^-/-^/Ca_v_3.1^-/-^ mutants did not change significantly (Table 1 and Supplementary Table 2). Together these data show that systemic injection of wild-type MDSCs into bradycardic mutant mice resulted in a significant improvement in the heart rate of mutant mice whereas it did not affect the heart rate of wild-type mice.

## Discussion

Our data reveal that skeletal muscle harbours a population of non-tumorigenic stem cells, MDSC, that can be directly transplanted by systemic injection and are capable of differentiating into functional pacemaker-like myocytes. Given the limitations of other sources that have been considered, MDSCs represent an attractive and easily available source of stem cells with a high potential for use in cell therapy for the treatment of disorders of human heart automaticity.

In the last decade, human myoblasts (derived from satellite cells) have been on preliminary clinical trial as a source of stem cells for myocardial autologous cell therapy raising hope of bringing functional benefit (reviewed by Menasché ^58^). However, it is now clearly recognized that myoblasts do not differentiate into cardiomyocytes after injection into the heart, but instead may fuse into multinucleated myotubes. Such differentiation may cause conduction blocks eventually leading to ventricular arrhythmia. This disappointing outcome essentially put a brake on considering skeletal muscle as a source of stem cells in cardiac cell therapy. Although most of the regenerative capacity of skeletal muscle throughout life is accounted for by satellite cells, it has become clear that the muscle niche is remodelled in dynamic balance with other stem cells derived from peri-vascular and peri-neural muscle compartment that represent additional sources of MDSCs ^2,9,10^.

Taking into account this alternative, our study opens a new avenue of research on the use of skeletal muscle as a source of stem cells in therapies for diseases of heart automaticity. Indeed, our data show that the systematic injection of undifferentiated MDSCs produced functional improvements in heart rhythm (and reduced arrhythmia) in recipient mutant mice, with IV- or IP-injected stem cells migrating and homing to the SAN region and differentiating into pacemaker-like cells expressing Ca_v_1.3 proteins. The current study is the first to demonstrate direct repair of the SAN through the *in situ* differentiation of transplanted stem cells. Alternative approaches towards effective biological pacing have been reported, with some using ion channel gene transfer in the ventricular conduction system ^22,59^ and others *in situ* reprogramming of ventricular cardiomyocytes ^60^. Although transplantation of a similar population of MDSCs has been reported to improve recovery of the heart from ischemic injury, the effect in that case was attributed to the secretion of paracrine factors without significant retrieval or differentiation of donor stem cells ^61^. ESCs and iPSCs can differentiate into beating, pacemaker-like cells *in vitro* ^47,48^. Such human ESC- and iPSC-derived pacemaker-like cells (1 to 2 × 10^6^ cells) were recently transplanted into rat left ventricle where they displayed pacemaker activity (for up to 60 second as measured on isolated Langerdorff-perfused hearts) despite less than 2% survival of the transplanted cells 2 weeks after transplantation^49,50^. In fact, the transforming and teratoma-forming potential of iPSC- and ESC-derived cells precludes their direct use for cell therapy and requires prior cueing toward cardiac differentiation, with a clear loss of plasticity, migration and survival ability ^47,62^. In contrast, MDSCs, which are non-tumorigenic, were directly transplanted into the bloodstream or peritoneal space of mutant recipient mice. After injection, MDSC were capable of (i) homing to the mutant SAN where they remained competent to divide for at least 72 hours, (ii) differentiating into arrays of Ca_v_1.3-expressing pacemaker-like cells in the SAN area (iii) electrically coupling with recipient SAN tissue to generate new leading pacemaker sites and (iv) reducing SAN dysfunction associated with the knock-out of the Ca_v_1.3 Ca^2+^ channel ^44^. The consistency between leading pacemaker sites in transplanted Ca_v_1.3^-/-^ mice and location of MDSC engraftment indicates that improvement of SAN function was not the result of paracrine activity of MDSC, but was due to electrical coupling between recipient SAN cells and differentiated MDSC-PM from donor mice. We also show here that MDSCs can differentiate *in vitro* into spontaneously active MDSC-PMs, displaying features and markers of mature SAN pacemaker cells and not those of atrial cells. SAN automaticity is generated by a functional association between the activity of ion membrane channels such as Ca_v_1.3 ^30^, HCN4 ^42^ and that of RyR-dependent Ca^2+^ release from the sarcoplasmic reticulum ^41^. Genetic ablation (from muscle-donor mouse) or pharmacologic inhibition of these channel activities in spontaneously active MDSC-PMs reproduced slowing of pacemaker activity, as reported in the current literature on native SAN-resident pacemaker cells. We found that some parameters (e.g. the kinetics of the repolarization phase) of the spontaneous action potential of *in vitro* differentiated MDSC-PM differ from those of native SAN myocytes. These discrepancies could be explained by the different conditions of spontaneous differentiation of MDSC-PM *in vitro*, or to a different degree of maturity of native SAN cells versus MDSC-PM. However, the similarity between the parameters of the pacemaker potential between the two cell types are comparable suggests common pacemaker mechanism(s). In addition, our results support the notion that MDSCs differentiated *in vitro* from mutant mice can reproduce and model dysfunctions of heart automaticity as confirmed when differentiating MDSC from Ca_V_1.3^-/-^ mouse muscle resulting into MDSC-PMs with slower pulsing rate (Fig.4e). This also may make MDSCs a convenient source of stem cells to recapitulate defects in heart automaticity *ex vivo* and assay potential therapeutic drugs. This hypothesis is in line with the capability of MDSCs to improve the heart rate of mutant mice following their colonization of the SAN and differentiation at this site.

Although the mechanism underlying the spontaneous ability of MDSCs to migrate and home to the SAN tissue of mutant mice is not fully understood, it likely involves chemotaxis via the SDF1-CXCR4 pathway, as supported by data shown in Supplementary Fig. 7. Furthermore, in light of reports on heart regeneration by exercise-activated cardiac stem cells ^63^, on the activation of muscle stem cells by exercise (review ^64^), and on the presence of stem cells from recipient patients in transplanted human hearts (i.e., “chimeric transplanted hearts”) ^65,66^, it is tempting to speculate that some stem cells that were classified as “resident cardiac” may have originated from exercise-activated circulating MDSCs. This hypothesis is further supported by recent reports on the absence of cardiac stem cells in adult heart. It is known that denervation, like exercise, results in long-lasting activation and proliferation of resident muscle stem cells but that 90% of these cells are subsequently lost from the muscle (and not fused to the denervated-re-innervated muscle) ^67^. In this regard, it will be important to compare profiles of circulating stem cells retrieved after exercise or muscle denervation to those in the resting state.

SAN dysfunction is a multi-factorial disease that often begins with a moderate reduction in heart rate that is otherwise asymptomatic and subsequently degenerates into bradycardia associated with invalidating symptoms and syncope ^68^. Early therapy of SAN dysfunction could potentially avoid or delay the need to implant an electronic pacemaker ^20^. In this regard, restoring damaged or degenerated SAN tissue by direct transplantation of MDSCs, in particular from the own muscle of the patient, constitutes an attractive possibility. We found that amelioration of heart rate in bradycardic mice occurs about 3 weeks following cell transplantation. This delay presumably reflects the time required for MDSCs to differentiate into MDSC-PMs. However, we recently also showed that the inhibition of G-protein gated K^+^ channels (*I*_*KACh*_) (by either genetic deletion or a pharmacologic agent) may rescue the SAN dysfunction and heart block normally observed in mice carrying silenced HCN4 channel conductance^33^ and Ca_v_1.3^-/-^ mice ^40^. It is thus possible that future therapies for SAN dysfunction may combine a pharmacologic strategy such as *I*_*KACh*_ inhibition to improve heart rate in the short term, followed by a long-term cell therapy-based approach employing MDSCs.

## Materials and Methods

### Mouse strains

Six to ten week-old wild-type C57Bl/6J mice or mTmG Tomato-expressing mice^52^ were used to prepare MDSC as donor mice for transplantation experiments. Recipient wild-type, Ca_v_1.3^-/-^ and Ca_v_1.3^-/-^/Ca_v_3.1^-/-^ mice used for transplantation experiments were all of C57Bl/6 genetic background ^57^. For transplantation experiments, we selected mice aged of at least 4 months of both sex to allow progression of the SAN dysfunction phenotype. All procedures were performed in accordance with the animal care guidelines of the European Union, French laws and the Ethical committee of the University of Montpellier (N°: 2017010310594939).

### Cell culture

MDSC were prepared using the preplate procedure originally described by Qu-Petersen et al.^11^ as modified in Arsic et al. ^10^ Briefly, mouse hind-limb muscles were removed, minced and enzymatically dissociated at 37°C for 45 min in 0.2% collagenase A (Roche) and 1 mg/ml dispase (GibcoBRL). After digestion, muscle cells were passed through 100 µm then 40 µm nylon cell strainers (BD Falcon) and centrifuged at 360 g for 10 min. Pelleted cells were washed twice in phosphate buffer saline (PBS), suspended in growth medium GM [DMEM/F12 (SIGMA, Les Ulis, France), 16% fetal bovine serum (PAA), 1% Ultroser G (Pall Life Sciences, Washington, USA), 100uM Beta-mercapto-ethanol, antibiotic-antimycotic mix (Gibco, Invitrogen Inc. Saint Quentin Fallavier, France) and plated on uncoated dishes (NUNC). After an initial 40 min pre-plating, non-adherent cells were collected and transferred to a new dish. The same operation was repeated every 24 h for 7 days with the addition of a centrifugation step at 360g, followed by resuspending the pelleted non-adherent cells in fresh GM.

MDSC used for RNA extraction, I.V. and I.P. transplantation and sub-cutaneous grafting experiments were non-adherent cells obtained after 5-7 days sequential pre-plating of stem cells extracted from hind limb skeletal muscle of adult mice.

Mouse adult fibroblast (MAF) were obtained from ATCC (Ltk-11) and grown as recommended by the supplier. Mouse ESCs were grown and harvested as described^69^.

### In vitro differentiation of MDSC

To characterize the pacemaker-like myocytes differentiated *in vitro* into autonomously beating cells (MDSC-PMs), MDSC were plated at a medium-high density (20-30,000 cells/cm2) on Polylysine- or fibronectin-coated dishes in DMEM/F12 containing 10% fetal bovine serum, 1% UltroserG (Pall Life Sciences, Washington USA) and antibiotic/antimycotic mix (Gibco/Invitrogen Inc). Under these conditions, MDSCs spontaneously differentiate into beating MDSC-PMs with the extent of differentiation reaching a maximal level by 3 to 4 weeks in culture and displaying a high density of contracting myocytes as shown in Supplementary video 3. (Video S3 shows one of the cultures of MDSC-PMs used in the gene expression profiling experiments at the time of harvesting the cells after 7 weeks in culture).

### Immunofluorescence analysis

Immunofluorescence analysis was performed on 2-6 week-cultured differentiated MDSC obtained from serial preplates as described^10^. Cells were fixed with 3.7% formaldehyde for 5 min, snap permeabilized for 30 sec in -20°C acetone, preincubated 30 min with 0.5% BSA in PBS. Cells were then incubated for 1h at 37°C with primary antibodies against HCN4, (1:100, Alomone), sarcomeric α-actinin, clone EA-53 (1:100, Sigma-Aldrich), connexin45 (1:100, H-85, Santa Cruz Biotechnology), Ca_v_1.3 polyclonal antibody^56^, cardiac troponin I (1:100, Millipore), Islet1 monoclonal antibody 39.3f7 (developed by Thomas M. Jessell and Susan Brenner-Morton and obtained from the Developmental Studies Hybridoma Bank, created by the NICHD of the NIH and maintained at The University of Iowa, Department of Biology, Iowa City, IA 52242). Specific staining was revealed using anti-mouse or anti-rabbit Alexa Fluor 488 and Alexa Fluor 555-conjugated secondary antibodies (InVitrogen) with Hoechst33358 nuclear stain (0.1 µg/ml) for 30 min at 37°C. The specificity of Ca_v_1.3 antibodies was demonstrated previously as the lack of immunostaining of SAN cells from Ca_v_1.3^-/-^ mice^56^ and verified in the present study (see Supplementary Fig 9). In the case of Cav1.3 antibodies, Ca_v_1.3^-/-^ SAN tissues served as controls.

### RNA isolation and reverse transcription polymerase chain reaction

Total RNA was extracted from sino-atrial node tissue, ventricular tissue, ESCs, mouse adult fibroblasts, MDSC and *in vitro* cultured differentiated MDSC using RNeasy kit (Qiagen) and then treated with RQ1 DNase treatement (Promega) according to the manufacturer’s recommendations. Ventricular and sino-atrial node tissues were obtained from 7-week-old C57Bl/6 adult male mice. Total RNA was used to synthesize cDNA using the Superscript III reverse transcriptase kit (Invitrogen) with random primers or with gene-specific primer in the case of Hcn4. The gene specific primers for Hcn4, were designed from alignment of four sequences (Hcn1, Hcn2, Hcn3 and Hcn4) and chosen on the basis of a specific N-terminal Hcn4 fragment. PCR was performed with GoTaq DNA polymerase (Promega); primer sequences are shown in supplemental Table S1. The cDNA was amplified using 35 cycles of PCR with an annealing temperature of 60°C to 62°C. PCR products were separated on 2% agarose gel (Invitrogen), stained with ethidium bromide; visualized and photographed on a UV transluminator (Fischer Scientific).

### Comparative RNAseq of MDSC, SAN, atria and MDSC-PM

For comparative RNA expression in whole muscle cells from first preplate: PP1 (at 24h) versus the 8^th^ preplate: PP8-MDSC shown Fig.1b: data shown are from pseudo-counts-RLE obtained after RNAseq analysis of triplicate samples from RNA banks made with the TruSeq Stranded mRNA sample preparation kit (Illumina) and sequencing with a HiSeq2500 Illumina sequencer (MGX Montpellier GenomiX: http://www.mgx.cnrs.fr). Data of RNAseq of native SAN, MDSC-PM and atrial tissue are expressed as pseudo-counts obtained after RNAseq analysis of samples from RNA banks (Supplementary Methods section).

### Cell transplantation experiments

Intravenous (I.V.) and intraperitoneal (I.P.) transplantation experiments were performed using wild-type, Ca_v_1.3^-/-^ or Ca_v_1.3^-/-^/Ca_v_3.1^-/-^ mice as recipients. MDSC were extracted from wild-type C57Bl/6J or mTmG Tomato-expressing mice^52^. Approximately 3 to 5×10^5^ cells per recipient mouse were resuspended in 0.9% sodium chloride solution and injected into the tail vein or into the peritoneal space. Mock-transplanted Ca_v_1.3^-/-^ mice were injected with sodium chloride solution alone. Transplanted mice were sacrificed 2, 10, 20, 30 and 41 days after injection followed by sino-atrial node dissection. For cell tracking experiments in the SAN tissue, MDSC were labelled before injection for 30 min with the lipophilic membrane-bound cell tracker DiI-CM (C7000 at 1µg/ml, ThermoFisher Scientific) according to the manufacturer’s instruction.

### Immunohistochemistry

Sino-atrial node tissue was dissected and fixed in 4% formaldehyde for 20 min before washing in PBS. After permeabilization in -20°C acetone, the whole SAN tissue was rehydrated with PBS. Non-specific binding of antibodies was blocked by incubation for 1h at room temperature with blocking solution (10% goat serum (Abcam), 0.1% Triton X-100 in PBS) followed by incubation with Mouse Ig Blocking Reagent (Vector Laboratories) for 30 min. Permeabilized and blocked sino-atrial tissue was then incubated overnight at 4°C with primary polyclonal anti-Ca_v_1.3-α1 subunit antibodies (prepared as described in ref. ^56^, 1:500) diluted in 1% goat serum in PBS, with mouse anti-Connexin45 (G-7, Santa Cruz) or with mouse anti-cdc6 antibody (37F4 from Molecular Probes). Alexa488- or 555-conjugated goat anti-rabbit and goat anti-mouse secondary antibodies (1:100, InVitrogen) in 1% goat serum were applied for 30 min at 37°C with Hoechst33358 nuclear stain (0.1 μg/ml) for 40 min at 37°C. Auto-fluorescence was quenched by incubating the tissue in 1% Sudan Black B (Sigma-Aldrich) in 70% ethanol for 10 min before washing in 70% ethanol for 5 min at room temperature. Tissues were mounted with AirVol mounting medium before photomicroscopy as described in Arsic et al.^10^. The specificity of Ca_v_1.3 staining was confirmed by its absence in sino-atrial tissue prepared from non-injected Ca_v_1.3^-/-^ mice.

### Measurements of contraction frequency

Cellular contraction of differentiated MDSC-PM was measured by an edge-detection IonOptix system device (LLC, 309 Hillside Street, Milton, MA 02186, USA). Cells were grown in 35 mm dishes in complete medium and analysed on an Olympus IX71 inverted microscope. MDSC-PM were imaged at 120 Hz using an IonOptix™ Myocam-S CCD camera. Digitized images were displayed within the IonWizard™ acquisition software (IonOptix™). To measure changes in cell length, two cursors were placed at opposing edges of the cell defined as difference in optical density (maximum: dark; minimum: light). The relative movement of cursors assessed cell shortening. Experiments were performed at 36 °C.

### Electrophysiological recordings of beating MDSC-PM, differentiated from MDSC in vitro

For electrophysiological recordings, aliquots of the cell suspension were harvested in custom made recording chambers (working volume 500 µL) allowing controlled unidirectional solution flow and mounted on the stage of an Olympus, X71 inverted microscope, and continuously perfused with normal Tyrode solution. The recording temperature was set to 36°C. The whole-cell variation of the patch-clamp technique^70^ was used to record cellular ionic currents, employing an Axopatch 200A (Axon Instruments Inc., Foster USA) patch-clamp amplifier, grounded by an agar bridge filled with 150 mM KCl. Cellular automaticity was recorded by the perforated patch technique with escin (50 μM). Recording electrodes were pulled from borosilicate glass, using a Flaming-Brown microelectrode puller (Sutter, Novato CA, USA). For recording cell automaticity, as well as *I*_*f*_ we used an intracellular solution containing (mM/L): K^+^-aspartate, 130; NaCl, 10; ATP-Na^+^ salt, 2; creatine phosphate, 6.6; GTP-Mg^2+^, 0.1; CaCl_2_, 0.04 (pCa=7); Hepes-KOH, 10; (adjusted to pH=7.2 with KOH). Electrodes had a resistance of about 5 MΩ. Seal resistances were in the range of 2-5 GΩ. The extracellular solution contained (in mM/L): NaCl, 140; KCl, 5.4; CaCl_2_, 1.8; MgCl_2_, 1; Hepes-NaOH, 5; and D-glucose, 5.5; (adjusted to pH=7.4 with NaOH). *I*_*f*_ was routinely recorded in Tyrode solution containing 5 mM BaCl_2_ to block *I*_*K1*_. Data was acquired with pClamp software (ver. 9, Axon Instruments Inc.).

### Confocal imaging of [Ca^2+^]_i_ transients and measurements of contraction

Intracellular Ca^2+^ ([Ca^2+^]_i_) transients were imaged by loading *in vitro* differentiated MDSC-PM with the Ca^2+^ indicator Fluo-4 AM 15 μM (from a stock solution in a mixture of DMSO/Pluronic F-127) in the medium for 45 min at 37°C. Spontaneous [Ca^2+^]_i_ transients were recorded in differentiated pulsing MDSC-PM loaded with Fluo-4 AM under control conditions (culture medium), or medium containing either epinephrine, acetylcholine (ACh) or ryanodine. Recordings were performed at 36°C. Differentiated MDSC-PM were distinguished from other cell types (i.e. fibroblasts and adipocytes cells) by their morphology (spindle and elongated shape), size (∼10 μm diameter) and pulsing activity. Extended Data videos S1-S3 show examples of differentiated MDSC-PM with typical morphology.

[Ca^2+^]_i_ transient images were obtained by confocal microscopy (Meta Zeiss LSM 510 and Zeiss LSM 780) scanning cells with an Argon laser in line scan configuration (1.54 ms line rate); fluorescence was excited at 488 nm and emissions collected at >505 nm. A 63x water immersion objective (n.a. 1.2), a 63x oil immersion objective (n.a. 1.32) and a 40x objective (n,a. 1.2) were used for recordings in differentiated pulsing MDSC-PM. Image analyses were performed using ImageJ software. Images were corrected for background fluorescence and the fluorescence values (F) normalized to basal fluorescence (F_0_) in order to obtain the fluorescence ratio (F/F_0_). Light intensity intervals were analyzed by pClamp software (ver. 9, Axon Instruments Inc.).

### Microscope and video imaging system

Digital images were collected using Leica DM LB microscope and Canon EOS300D digital camera as described before^10^. Videos of contracting cells were recorded using a NIKON D90 camera coupled to a Zeiss Axiovert 35M. A Nanozoomer Hamamatsu blade scanner was used for 2D reconstruction of whole mount SAN tissue (in Fig. S5 and Fig. S6).

### Tumour formation assay

1×10^6^ MDSC resuspended in 0.2 ml of 1/2X matrigel in HBSS were injected subcutaneously into the back of n=4 8-week-old SCID/beige mice (Charles River). Follow up observation was carried out for up to 4 months. ESCs derived tumours were obtained by similar subcutaneous injection of 1×10^6^ mouse ESCs into the back of 8-week-old SCID/beige mice (n=2). Tumours developed in all ES-grafted mice within 2 weeks. The mice were euthanized after 4 weeks when tumour size reached more than 2 cm.

### Optical mapping of membrane voltage in SAN/atria preparations

To analyse changes in membrane voltage in SAN/atrial tissue, the entire preparation, including the SAN, the left and the right atrium, was loaded by immersing the tissue in a Tyrode’s solution containing the voltage-sensitive indicator/dye Di-4-ANEPPS (10 µM; Biotium) for at least 30 min at room temperature (20–22 °C). The preparation was placed on agitated plate during loading to maintain proper oxygenation of the tissue and load it uniformly. After the loading step, the tissue was washed 3 times in dye-free Tyrode’s solution. The SAN/atrial tissue was then constantly perfused at 34–36 °C and imaged by high speed optical voltage mapping (2 ms per frame) on a MiCAM Ultima-L complementary metal oxide semiconductor (CMOS) camera (100×100-pixel CMOS sensor, 10×10 mm, SciMedia). This camera was mounted on a THT microscope, with two objectives (2×and 1.6×) that generated a field of view of 12.5×12.5 mm. A 150-W halogen light system with built-in shutter (SciMedia) was used as an excitation light source for the voltage dye. The filter set included a 531/50-nm excitation filter, 580-nm dichroic mirror, and 580 long-pass emission filter. To avoid motion artifacts, we blocked mechanical activity using blebbistatin (1.5–5µM; Tocris Bioscience). We usually limited our recording times to 32.768 s (16,384 frames at 2 ms per frame) to avoid phototoxic effects of the dye. Optical raw data were analyzed using dedicated software from the camera developer, BVAna Analysis Software (Brainvision).

### ECG recordings and analysis in sedated mice

One-lead surface ECG measurements were recorded from wild-type and Ca_v_1.3^-/-^/Ca_v_3.1^-/-^ mice sedated with 1.5% isofluorane. Body temperature was continuously maintained at 36° -37°C using a heated pad connected to a controller that received feedback from a temperature sensor attached to the mouse. Ag/AgCl gel-coated ECG electrodes (Unomedical) were attached to the superior right and both inferior limbs of mice. The electrodes were connected to a standard one-lead ECG amplifier module (EMKA Technologies, Paris, France), which included high- and low-pass filters (set to 0.05 Hz and 500 Hz, respectively) and a gain selection device (set to 1,000-fold). Signals were digitized continuously at 2 kHz and recorded using an IOX data acquisition system (EMKA Technologies, France). The recordings were carried out for a 45-min period before (day 0) and at different days after transplantation (day 10 and day 20). The software ecgAuto (EMKA Technologies, France) was used to perform offline analysis of the data recorded. For each mouse the mean heart rate value and the Standard Deviation (SD) were calculated. Beat-to-beat analysis of heart rate was carried out on a 30-min interval taken 10 min after the beginning of each 45-min recording.

### Telemetric recordings of ECG and analysis

For telemetric ECG recording, adult wild-type and Ca_v_1.3^-/-^ male mice were anesthetized with 2% isofluorane. A midline incision was made on the back along the spine to insert a telemetric transmitter (TA10EA-F20, Data Sciences International) into a subcutaneous pocket with paired wire electrodes placed over the thorax (chest bipolar ECG lead). Local anaesthesia was obtained with lidocaine (1%) injected subcutaneously at the sites of electrodes and transmitter implantation. To manage possible post-surgery pain, Advil (paracetamol and ibuprofene, 7 mL/l) was added to the drinking water for four days after implantation. Mice were housed in individual cages with free access to food and water and were exposed to 24-hour light/dark cycles (light, 8:30 AM to 8:30 AM) in a thermostatically controlled room. Heart rates were averaged over a 24-hour recording period or over two 12-hour periods corresponding to “day-time” (light, 8:30 AM to 8.30 PM) or “night-time” (dark, 8.30 PM to 8.30 AM). Recordings were initiated at least 8 days after recovery from surgical implantation. ECG signals were recorded using a telemetry receiver and an analog-to-digital conversion data acquisition system for display and analysis by DataquestTM A.R.T.TM software (Data Sciences International). ECG parameters were measured using ecgAuto 1.5.7 software. RR, PR, QRS, QT and QTc intervals were first calculated automatically by ecgAuto based on a large (>250) ECG sample library and then verified by visual inspection. Heart rates (in bpm) were then determined from RR intervals. PP intervals were measured manually over shorter recording windows.

### Statistical analysis

The statistical significance was evaluated through appropriated statistical tests, as reported in each figure legend. All experimental samples tested and all animals recruited in the experimental plan were included in data analysis, to ensure maximal representativity in data sections. No method of blinding to the experimenter was adopted in data recordings and analysis. Statistical significance was set at p<0.05. Data analysis was performed with GraphPad Prism 6.0.

### Data availability

The data that support the findings of this study are available from the authors on reasonable request, see author contributions for specific data sets.

## Supporting information

Supplementary-data-and-figures-Rev1

## Author Contributions

A.F. and M.E.M. designed and analysed the experiments. P.M., D.M., I.B. performed the experiments with additional input from M.B., A.G.T., V.M., R.D., M.L.D.F., N.A., J.N., A.L., J.S. and N.J.C.; D.M. and R.D. performed muscle stem cell culture and differentiation, RT-PCR analysis with advice from N.A. and immunocyto- and histochemistry with supervision from A.F. and N.J.L.; P.M. performed whole-cell patch clamp, Ca^2+^ imaging and ECG telemetric recordings with supervision from M.E.M.; M.B. performed optical mapping experiments with the supervision of P.M. and M.E.M.; A.F. and M.E.M wrote the manuscript with comments and approval from all co-authors.

## Acknowledgements

We thank Xavier Hautecoeur (IGH, Montpellier) for help with MDSC culture and Chantal Jacquet (IGMM, Montpellier) for initial help with MDSC I.V. injection. We are grateful to the personnel of the MRI, RHEM, RAM and MGX facilities of Biocampus Montpellier. We thank Dr. Olivier Ganier (IGH) for mouse embryonic stem cells. The hybridoma monoclonal antibodies Islet1 39.3f7 developed by Thomas M. Jessell and Susan Brenner-Morton was obtained from the Developmental Studies Hybridoma Bank, created by the NICHD of the NIH and maintained at The University of Iowa, Department of Biology, Iowa City, IA 52242.

## Funding

P.M. and A.G.T. were supported by a Research Training Network (RTN) “CavNet” funded through the EU Research Programme (6FP) MRTN-CT-2006-035367. The IGF group is a member of the Laboratory of Excellence (LabEx) “Ion Channel Science and Therapeutics” (ICST) supported by a grant from ANR (ANR-11-LABX-0015). M.B. was supported by a LabEx ICST Ph.D. fellowship. D.M. and V.M. were supported by Association Francaise contre les Myopathies (AFM). Research supported by the Fondation pour la Recherche Medicale “Physiopathologie Cardiovasculaire” (DPC20111122986 to A.F. and DPC20171138970 to M.M.), by Agence Nationale de Recherche (ANR-2010-BLAN-1128-01 to MEM/AF and ANR-15-CE14-0004-01 to M.M.), NIH (DC009433, HL087120 to A.L.), University of Iowa Carver Research Program of Excellence Award (to A.L.), the Austrian Science Fund (F27809 to J.S.).

## Notes

#### Summary of Updates

This revised manuscript is as submitted to Nat. Comm in summer 2019 and a further resubmission is pending. The manuscript has been revised to provide further data that: - increased expression of characteristic markers of pacemaker cells occurs during in vitro differentiation of MDSC. - in vivo migration and homing of MDSC to the Sino-atrial node (SAN) of bradycardic mutant mice takes place and involves increased levels of the chemokine SDF1 and its receptor CXCR4 in mutant SAN tissue. - MDSC integration as pacemaking cells occurs in mutant recipient mouse SAN as detected both electrically and functionally.

## References

1 Peault, B. et al. Stem and progenitor cells in skeletal muscle development, maintenance, and therapy. Mol Ther 15, 867–877, doi:6300145 [pii] 10.1038/mt.sj.6300145 (2007).

2 Tamaki, T. Bridging long gap peripheral nerve injury using skeletal muscle-derived multipotent stem cells. Neural Regen Res 9, 1333–1336, doi:10.4103/1673-5374.137582 (2014).

3 Tamaki, T., Uchiyama, Y. & Akatsuka, A. Plasticity and physiological role of stem cells derived from skeletal muscle interstitium: contribution to muscle fiber hyperplasia and therapeutic use. Curr Pharm Des 16, 956–967 (2010).

4 Yin, H., Price, F. & Rudnicki, M. A. Satellite cells and the muscle stem cell niche. Physiol Rev 93, 23–67, doi:10.1152/physrev.00043.2011 (2013).

5 Ten Broek, R. W., Grefte, S. & Von den Hoff, J. W. Regulatory factors and cell populations involved in skeletal muscle regeneration. J Cell Physiol 224, 7–16, doi:10.1002/jcp.22127 (2010).

6 Romero-Ramos, M. et al. Neuronal differentiation of stem cells isolated from adult muscle. J Neurosci Res 69, 894–907, doi:10.1002/jnr.10374 (2002).

7 Alessandri, G. et al. Isolation and culture of human muscle-derived stem cells able to differentiate into myogenic and neurogenic cell lineages. Lancet 364, 1872–1883, doi:S0140-6736(04)17443-6 [pii] 10.1016/S0140-6736(04)17443-6 (2004).

8 Uezumi, A. et al. Functional heterogeneity of side population cells in skeletal muscle. Biochem Biophys Res Commun 341, 864–873, doi:S0006-291X(06)00104-5 [pii] 10.1016/j.bbrc.2006.01.037 (2006).

9 Tamaki, T. et al. Clonal multipotency of skeletal muscle-derived stem cells between mesodermal and ectodermal lineage. Stem Cells 25, 2283–2290, doi:10.1634/stemcells.2006-0746 (2007).

10 Arsic, N., Mamaeva, D., Lamb, N. J. & Fernandez, A. Muscle-derived stem cells isolated as non-adherent population give rise to cardiac, skeletal muscle and neural lineages. Exp Cell Res 314, 1266–1280, doi:S0014-4827(08)00019-0 [pii] 10.1016/j.yexcr.2008.01.009 (2008).

11 Qu-Petersen, Z. et al. Identification of a novel population of muscle stem cells in mice: potential for muscle regeneration. J Cell Biol 157, 851–864, doi:10.1083/jcb.200108150 jcb.200108150 [pii] (2002).

12 Gharaibeh, B. et al. Isolation of a slowly adhering cell fraction containing stem cells from murine skeletal muscle by the preplate technique. Nat Protoc 3, 1501–1509, doi:nprot.2008.142 [pii] 10.1038/nprot.2008.142 (2008).

13 Mangoni, M. E. & Nargeot, J. Genesis and regulation of the heart automaticity. Physiol Rev 88, 919–982 (2008).

14 Christoffels, V. M., Smits, G. J., Kispert, A. & Moorman, A. F. Development of the pacemaker tissues of the heart. Circ Res 106, 240–254, doi:106/2/240 [pii] 10.1161/CIRCRESAHA.109.205419 (2010).

15 Wiese, C. et al. Formation of the sinus node head and differentiation of sinus node myocardium are independently regulated by Tbx18 and Tbx3. Circ Res 104, 388–397, doi:CIRCRESAHA.108.187062 [pii] 10.1161/CIRCRESAHA.108.187062 (2009).

16 Verheijck, E. E., Wilders, R. & Bouman, L. N. Atrio-sinus interaction demonstrated by blockade of the rapid delayed rectifier current. Circulation 105, 880–885 (2002).

17 Watanabe, E. I. et al. Modulation of pacemaker activity of sinoatrial node cells by electrical load imposed by an atrial cell model. Am J Physiol 269, H1735–1742 (1995).

18 Verheule, S. & Kaese, S. Connexin diversity in the heart: insights from transgenic mouse models. Front Pharmacol 4, 81, doi:10.3389/fphar.2013.00081 (2013).

19 Mesirca, P., Torrente, A. G. & Mangoni, M. E. Functional role of voltage gated Ca(2+) channels in heart automaticity. Front Physiol 6, 19, doi:10.3389/fphys.2015.00019 (2015).

20 Jensen, P. N. et al. Incidence of and risk factors for sick sinus syndrome in the general population. J Am Coll Cardiol 64, 531–538, doi:S0735-1097(14)02843-5 [pii] 10.1016/j.jacc.2014.03.056 (2014).

21 Mond, H. G. & Proclemer, A. The 11th world survey of cardiac pacing and implantable cardioverter-defibrillators: calendar year 2009--a World Society of Arrhythmia’s project. Pacing Clin Electrophysiol 34, 1013–1027, doi:10.1111/j.1540-8159.2011.03150.x (2011).

22 Rosen, M. R., Robinson, R. B., Brink, P. R. & Cohen, I. S. The road to biological pacing. Nat Rev Cardiol 8, 656–666, doi:nrcardio.2011.120 [pii] 10.1038/nrcardio.2011.120 (2011).

23 Besson, V. et al. PW1 gene/paternally expressed gene 3 (PW1/Peg3) identifies multiple adult stem and progenitor cell populations. Proc Natl Acad Sci U S A 108, 11470–11475, doi:10.1073/pnas.1103873108 (2011).

24 Hoogaars, W. M. et al. Tbx3 controls the sinoatrial node gene program and imposes pacemaker function on the atria. Genes Dev 21, 1098–1112 (2007).

25 Ye, W. et al. A common Shox2-Nkx2-5 antagonistic mechanism primes the pacemaker cell fate in the pulmonary vein myocardium and sinoatrial node. Development 142, 2521–2532, doi:10.1242/dev.120220 (2015).

26 Hashem, S. I. et al. Shox2 regulates the pacemaker gene program in embryoid bodies. Stem Cells Dev 22, 2915–2926, doi:10.1089/scd.2013.0123 (2013).

27 Weinberger, F. et al. Localization of Islet-1-positive cells in the healthy and infarcted adult murine heart. Circ Res 110, 1303–1310, doi:CIRCRESAHA.111.259630 [pii] 10.1161/CIRCRESAHA.111.259630 (2012).

28 Ludwig, A., Zong, X., Jeglitsch, M., Hofmann, F. & Biel, M. A family of hyperpolarization-activated mammalian cation channels. Nature 393, 587–591 (1998).

29 Brown, H. F., DiFrancesco, D. & Noble, S. J. How does adrenaline accelerate the heart? Nature 280, 235–236 (1979).

30 Mangoni, M. E. et al. Functional role of L-type Cav1.3 Ca2+ channels in cardiac pacemaker activity. Proc Natl Acad Sci U S A 100, 5543–5548 (2003).

31 Alig, J. et al. Control of heart rate by cAMP sensitivity of HCN channels. Proc Natl Acad Sci U S A 106, 12189–12194 (2009).

32 Baruscotti, M. et al. Deep bradycardia and heart block caused by inducible cardiac-specific knockout of the pacemaker channel gene Hcn4. Proc Natl Acad Sci U S A 108, 1705–1710, doi:1010122108 [pii] 10.1073/pnas.1010122108 (2011).

33 Mesirca, P. et al. Cardiac arrhythmia induced by genetic silencing of ‘funny’ (f) channels is rescued by GIRK4 inactivation. Nat Commun 5, 4664, doi:ncomms5664 [pii] 10.1038/ncomms5664 (2014).

34 Torrente, A. G. et al. L-type Cav1.3 channels regulate ryanodine receptor-dependent Ca2+ release during sino-atrial node pacemaker activity. Cardiovasc Res 109, 451–461, doi:cvw006 [pii] 10.1093/cvr/cvw006 (2016).

35 Milanesi, R., Baruscotti, M., Gnecchi-Ruscone, T. & DiFrancesco, D. Familial sinus bradycardia associated with a mutation in the cardiac pacemaker channel. N Engl J Med 354, 151–157 (2006).

36 Baig, S. M. et al. Loss of Ca(v)1.3 (CACNA1D) function in a human channelopathy with bradycardia and congenital deafness. Nat Neurosci 14, 77–84, doi:nn.2694 [pii] 10.1038/nn.2694 (2011).

37 Verheijck, E. E. et al. Electrophysiological features of the mouse sinoatrial node in relation to connexin distribution. Cardiovasc Res 52, 40–50 (2001).

38 Vedantham, V., Galang, G., Evangelista, M., Deo, R. C. & Srivastava, D. RNA sequencing of mouse sinoatrial node reveals an upstream regulatory role for Islet-1 in cardiac pacemaker cells. Circ Res 116, 797–803, doi:10.1161/CIRCRESAHA.116.305913 (2015).

39 Linscheid, N. et al. Quantitative proteomics and single-nucleus transcriptomics of the sinus node elucidates the foundation of cardiac pacemaking. Nat Commun 10, 2889, doi:10.1038/s41467-019-10709-9 (2019).

40 Mesirca, P. et al. G protein-gated IKACh channels as therapeutic targets for treatment of sick sinus syndrome and heart block. Proc Natl Acad Sci U S A 113, E932–941, doi:1517181113 [pii] 10.1073/pnas.1517181113 (2016).

41 Lakatta, E. G., Maltsev, V. A. & Vinogradova, T. M. A coupled SYSTEM of intracellular Ca2+ clocks and surface membrane voltage clocks controls the timekeeping mechanism of the heart’s pacemaker. Circ Res 106, 659–673, doi:106/4/659 [pii] 10.1161/CIRCRESAHA.109.206078 (2010).

42 DiFrancesco, D. The role of the funny current in pacemaker activity. Circ Res 106, 434–446, doi:106/3/434 [pii] 10.1161/CIRCRESAHA.109.208041 (2010).

43 Bois, P., Bescond, J., Renaudon, B. & Lenfant, J. Mode of action of bradycardic agent, S 16257, on ionic currents of rabbit sinoatrial node cells. Br J Pharmacol 118, 1051–1057 (1996).

44 Platzer, J. et al. Congenital deafness and sinoatrial node dysfunction in mice lacking class D L-type Ca2+ channels. Cell 102, 89–97 (2000).

45 Zhang, Z. et al. Functional Roles of Ca(v)1.3 (alpha(1D)) calcium channel in sinoatrial nodes: insight gained using gene-targeted null mutant mice. Circ Res 90, 981–987. (2002).

46 Liaqat, K. et al. Identification of CACNA1D variants associated with sinoatrial node dysfunction and deafness in additional Pakistani families reveals a clinical significance. J Hum Genet 64, 153–160, doi:10.1038/s10038-018-0542-8 (2019).

47 Laflamme, M. A. & Murry, C. E. Heart regeneration. Nature 473, 326–335, doi:nature10147 [pii] 10.1038/nature10147 (2011).

48 Kleger, A. et al. Modulation of calcium-activated potassium channels induces cardiogenesis of pluripotent stem cells and enrichment of pacemaker-like cells. Circulation 122, 1823–1836, doi:CIRCULATIONAHA.110.971721 [pii] 10.1161/CIRCULATIONAHA.110.971721 (2010).

49 Chauveau, S. et al. Induced Pluripotent Stem Cell-Derived Cardiomyocytes Provide In Vivo Biological Pacemaker Function. Circ Arrhythm Electrophysiol 10, e004508, doi:CIRCEP.116.004508 [pii] 10.1161/CIRCEP.116.004508 (2017).

50 Protze, S. I. et al. Sinoatrial node cardiomyocytes derived from human pluripotent cells function as a biological pacemaker. Nat Biotechnol 35, 56–68, doi:10.1038/nbt.3745 (2017).

51 Mitutsova, V. et al. Adult muscle-derived stem cells engraft and differentiate into insulin-expressing cells in pancreatic islets of diabetic mice. Stem Cell Res Ther 8, 86, doi:10.1186/s13287-017-0539-9 (2017).

52 Muzumdar, M. D., Tasic, B., Miyamichi, K., Li, L. & Luo, L. A global double-fluorescent Cre reporter mouse. Genesis 45, 593–605, doi:10.1002/dvg.20335 (2007).

53 Hateboer, G. et al. Cell cycle-regulated expression of mammalian CDC6 is dependent on E2F. Mol Cell Biol 18, 6679–6697, doi:10.1128/mcb.18.11.6679 (1998).

54 Miller, R. J., Banisadr, G. & Bhattacharyya, B. J. CXCR4 signaling in the regulation of stem cell migration and development. J Neuroimmunol 198, 31–38, doi:10.1016/j.jneuroim.2008.04.008 (2008).

55 Stumm, R. & Hollt, V. CXC chemokine receptor 4 regulates neuronal migration and axonal pathfinding in the developing nervous system: implications for neuronal regeneration in the adult brain. J Mol Endocrinol 38, 377–382, doi:10.1677/JME-06-0032 (2007).

56 Christel, C. J. et al. Distinct localization and modulation of Cav1.2 and Cav1.3 L-type Ca2+ channels in mouse sinoatrial node. J Physiol 590, 6327–6342, doi:jphysiol.2012.239954 [pii] 10.1113/jphysiol.2012.239954 (2012).

57 Marger, L. et al. Functional roles of Ca(v)1.3, Ca(v)3.1 and HCN channels in automaticity of mouse atrioventricular cells: insights into the atrioventricular pacemaker mechanism. Channels (Austin) 5, 251–261, doi:15266 [pii] (2011).

58 Menasche, P. Stem cell therapy for chronic heart failure: lessons from a 15-year experience. C R Biol 334, 489–496, doi:10.1016/j.crvi.2011.03.006 (2011).

59 Boink, G. J. et al. HCN2/SkM1 gene transfer into canine left bundle branch induces stable, autonomically responsive biological pacing at physiological heart rates. J Am Coll Cardiol 61, 1192–1201, doi:S0735-1097(13)00182-4 [pii] 10.1016/j.jacc.2012.12.031 (2013).

60 Kapoor, N., Liang, W., Marban, E. & Cho, H. C. Direct conversion of quiescent cardiomyocytes to pacemaker cells by expression of Tbx18. Nat Biotechnol 31, 54–62, doi:10.1038/nbt.2465 (2013).

61 Okada, M. et al. Human skeletal muscle cells with a slow adhesion rate after isolation and an enhanced stress resistance improve function of ischemic hearts. Mol Ther 20, 138–145, doi:10.1038/mt.2011.229 (2012).

62 Mohsin, S., Siddiqi, S., Collins, B. & Sussman, M. A. Empowering adult stem cells for myocardial regeneration. Circ Res 109, 1415–1428, doi:109/12/1415 [pii] 10.1161/CIRCRESAHA.111.243071 (2011).

63 Ellison, G. M. et al. Adult c-kit(pos) cardiac stem cells are necessary and sufficient for functional cardiac regeneration and repair. Cell 154, 827–842 (2013).

64 Macaluso, F. & Myburgh, K. H. Current evidence that exercise can increase the number of adult stem cells. J Muscle Res Cell Motil 33, 187–198, doi:10.1007/s10974-012-9302-0 (2012).

65 Quaini, F. et al. Chimerism of the transplanted heart. N Engl J Med 346, 5–15, doi:10.1056/NEJMoa012081 (2002).

66 Laflamme, M. A., Myerson, D., Saffitz, J. E. & Murry, C. E. Evidence for cardiomyocyte repopulation by extracardiac progenitors in transplanted human hearts. Circ Res 90, 634–640 (2002).

67 Murray, M. A. & Robbins, N. Cell proliferation in denervated muscle: time course, distribution and relation to disuse. Neuroscience 7, 1817–1822 (1982).

68 Semelka, M., Gera, J. & Usman, S. Sick sinus syndrome: a review. Am Fam Physician 87, 691–696, doi:d10507 [pii] (2013).

69 Ganier, O. et al. Synergic reprogramming of mammalian cells by combined exposure to mitotic Xenopus egg extracts and transcription factors. Proc Natl Acad Sci U S A 108, 17331–17336, doi:1100733108 [pii] 10.1073/pnas.1100733108 (2011).

70 Hamill, O. P., Marty, A., Neher, E., Sakmann, B. & Sigworth, F. J. Improved patch-clamp techniques for high-resolution current recording from cells and cell-free membrane patches. Pflugers Archiv - European Journal of Physiology 391, 85–100 (1981).

