## Supplementary-data-and-figures-Rev1 for "Bradycardic mice undergo effective heart rate improvement after specific homing to the sino-atrial node and differentiation of adult muscle derived stem cells"

### Supplementary Methods

RNA Sequencing

Poly-A based mRNA enrichment and reverse transcription: RNA-Seq libraries were constructed with the Truseq stranded mRNA sample preparation (Low throughput protocol) kit from Illumina.

Five hundred nanogram of total RNA was used for the construction of the libraries. The first step in the workflow involves purifying the poly-A containing mRNA molecules using poly-T oligo attached magnetic beads. Following purification, the mRNA is fragmented into small pieces using divalent cations under elevated temperature. The cleaved RNA fragments are copied into first strand cDNA using SuperScript II reverse transcriptase, Actinomycin D and random hexamer primers. The Second strand cDNA was synthesized by replacing dTTP with dUTP. These cDNA fragments then have the addition of a single 'A' base and subsequent ligation of the adapter. The products are then purified and enriched with 15 cycles of PCR. The final cDNA libraries were validated with a Fragment Analyzer system (Advanced Analytical Technologies, Ankeny, IA) and quantified with a KAPA qPCR kit.

Sequencing: For each sequencing lane of a flowcell V4, three libraries were pooled in equal proportions, denatured with NaOH and diluted to 22 pM before clustering. Cluster formation, primer hybridisation and single end-read 50 cycles sequencing were performed on cBot and HiSeq2500 (Illumina, San Diego, CA) respectively.

Sequencing quality control: Image analyses and base calling were performed using the Illumina HiSeq Control Software and the Real-Time Analysis component. Demultiplexing was performed using Illumina's conversion software (bcl2fastq 2.20). The quality of the raw data was assessed using FastQC from the Babraham Institute and the Illumina software SAV (Sequencing Analysis Viewer). Potential contaminants were monitored with the FastQ Screen software from the Babraham Institute.

RNA-Seq data analysis: We aligned RNA-seq reads to the mouse genome (UCSC mm10) with the splice junction mapper TopHat 2.1.1[1], which used Bowtie 2.3.4.3[2]. We downloaded gene model annotations from the UCSC database (genes.gtf 15 January 2019). Final read alignments having more than 3 mismatches were discarded. We performed gene counting using featureCounts 1.6.2[3]. Counts were normalized using the trimmed mean of M-values (TMM) method implemented in the Bioconductor[4] package edgeR 3.20.1[5].

### References

1. Kim, D. *et al.* TopHat2: accurate alignment of transcriptomes in the presence of insertions, deletions and gene fusions. *Genome Biol* **14**, R36, doi:10.1186/gb-2013-14-4-r36 (2013).
2. Langmead, B. & Salzberg, S. L. Fast gapped-read alignment with Bowtie 2. *Nat Methods* **9**, 357-359, doi:10.1038/nmeth.1923 (2012).
3. Liao, Y., Smyth, G. K. & Shi, W. featureCounts: an efficient general purpose program for assigning sequence reads to genomic features. *Bioinformatics* **30**, 923-930, doi:10.1093/bioinformatics/btt656 (2014).
4. Gentleman, R. C. *et al.* Bioconductor: open software development for computational biology and bioinformatics. *Genome Biol* **5**, R80, doi:10.1186/gb-2004-5-10-r80 (2004).
5. Robinson, M. D., McCarthy, D. J. & Smyth, G. K. edgeR: a Bioconductor package for differential expression analysis of digital gene expression data. *Bioinformatics* **26**, 139-140, doi:10.1093/bioinformatics/btp616 (2010).

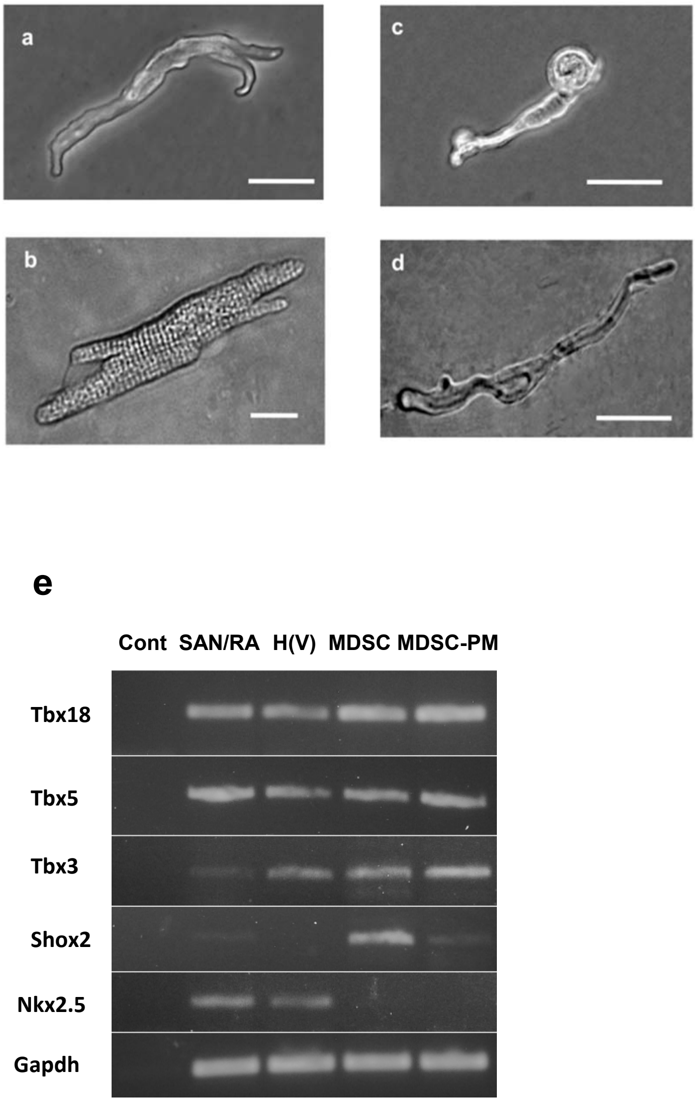


**Supplementary Fig. 1.**
**Morphological features of native pacemaker cells, atrial cells and MDSC-PMs.** Phase-contrast images of (**a**) a native pacemaker cell isolated from adult mouse SAN and (**b**) atrial cells isolated from adult mouse right atrium (**c,d**). Two examples of isolated beating MDSC-PMs differentiated *in vitro* from cultured MDSCs, following enzymatic detachment for patch-clamp analysis (shown in Fig. 4). Bars, 10 µm
(**e**)**. RT-PCR-based analysis of expression of transcription factors involved in sino-atrial node (SAN) development** (Tbx18, Tbx5, Tbx3, Shox2 and Nkx2.5). SAN/RA: mouse sino-atrial node and right atrium tissue; H(V): heart ventricles; MDSCs: muscle-derived stem cell; MDSC-PMs: MDSCs differentiated *in vitro* into pacemaker-like beating cells. Gapdh was used as loading control.

**Supplementary Table 1.** PCR primers used

| Gene | Primer | Product size, bp | Accession number |
|---|---|---|---|
| Hcn4 | 5'-GACGAGGAAGAGGATGGTGA-3'<br>5'-CACCGTTGGTGCTGGACT-3' | 196 | NM_001081192.1 |
| Nkx2-5 | 5'-CAAGTGCTCTCCTGCTTTCC-3'<br>5'-CGGCTTTGTCCAGCTCCACT-3' | 137 | NM_008700.2 |
| Isl1 | 5'-CGACCCAGTCAATGGAAACT-3'<br>5'-TGGGCTTAGGGTTTTGTTTG-3' | 155 | NM_021459.4 |
| Tbx18 | 5'-GGGGAGACTTGGATGAGACA-3'<br>5'-GGACAGATCATCTCCGCAAT-3' | 153 | NM_023814.4 |
| Tbx3 | 5'-GCTGCTGCGAACTCTCTTCT-3'<br>5'-ACCAATTGTGTGGCTGCATA-3' | 233 | NM_011535.2 |
| Tnni3 | 5’-TAAGATCTCCGCCTCCAGAA-3’<br>5’-CGGCATAAGTCCTGAAGCTC-3’ | 177 | NM_009406.3 |
| Cacna1D<br>($Ca_v1.3$) | 5’-TGCACAGATGAAGCCAAAAG-3’<br>5’-GACCAACGTTCTCACCGTTT-3’ | 229 | NM_001083616.1<br>NM_028981.2 |
| Actc1 | 5’-CTGAGATGTCTCTCTCTTAGCCTA-3’<br>5’-ACAATGACTGATGAGAGATG-3’ | 99 | NM_009608.3 |
| Cx45 | 5’-TGGTTGGGCTTAAAACTTGG-3’<br>5’-CAGCTCCACCTTCAGAGTCC-3’ | 151 | NM_001159382.1 |
| Shox2 | 5'-CCCTTGTCCTTTCAGGTTCA-3'<br>5'-ATGCTGGAGTTCTTGCTGGT-3' | 221 | NM_013665.1 |
| Gapdh | 5'-ACCCAGAAGACTGTGGATGG-3'<br>5'-CACATTGGGGGTAGGAACAC-3' | 171 | NM_008084.2 |

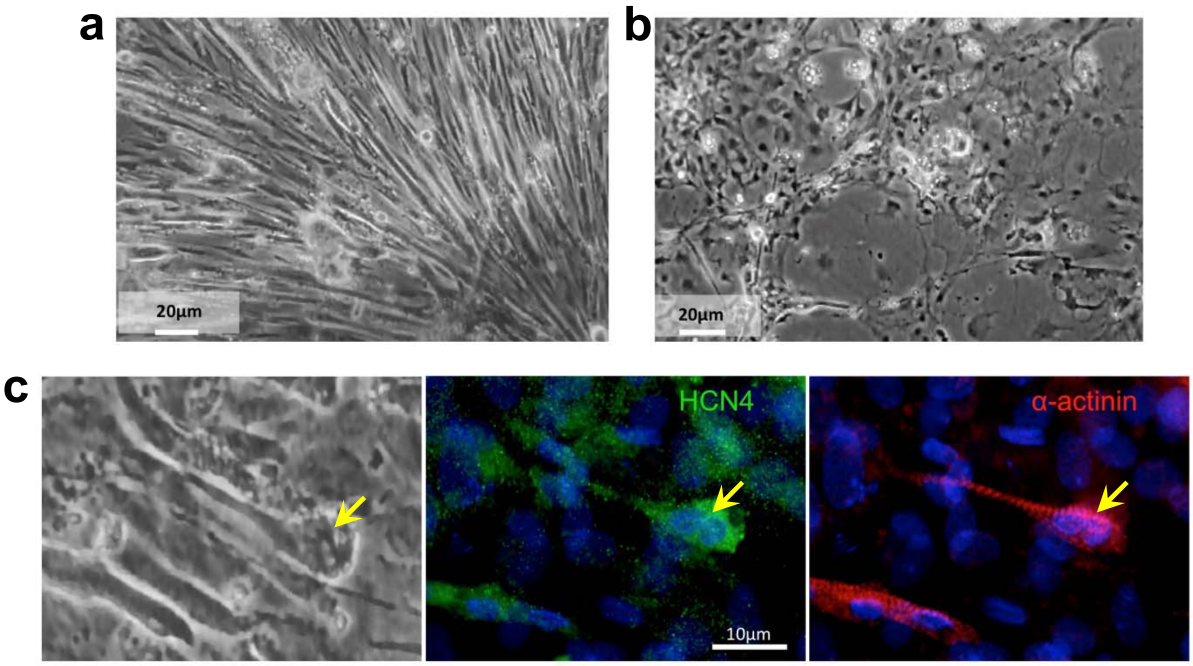


**Supplementary Fig. 2: *In vitro* differentiation of rat-muscle MDSCs into MDSC-PMs and co-staining for specific pacemaker markers. a** Field of cultured MDSCs at early stages of differentiation, 8 days after completing the isolation of MDSC from rat hind-limb muscle. Several cells in this field exhibited sustained and rhythmic contraction whereas others did not . **b** Field of cultured MDSCs at a later stages of differentiation, 36 days after isolation of MDSCs. In this culture, a small number of small cells contracted rhythmically amongst fibroblasts and adipocytes. **c.** Close up field from differentiated MDSCs in **b.** showing phase contrast (left), immunostaining for HCN4 (middle) and for alpha-actinin (right). Video recording of the live cells in **c.** is shown in Supplementary video 5. Arrow indicates a single bi-nucleated, contracting cell that co-expresses HCN4 and $\alpha$-actinin.

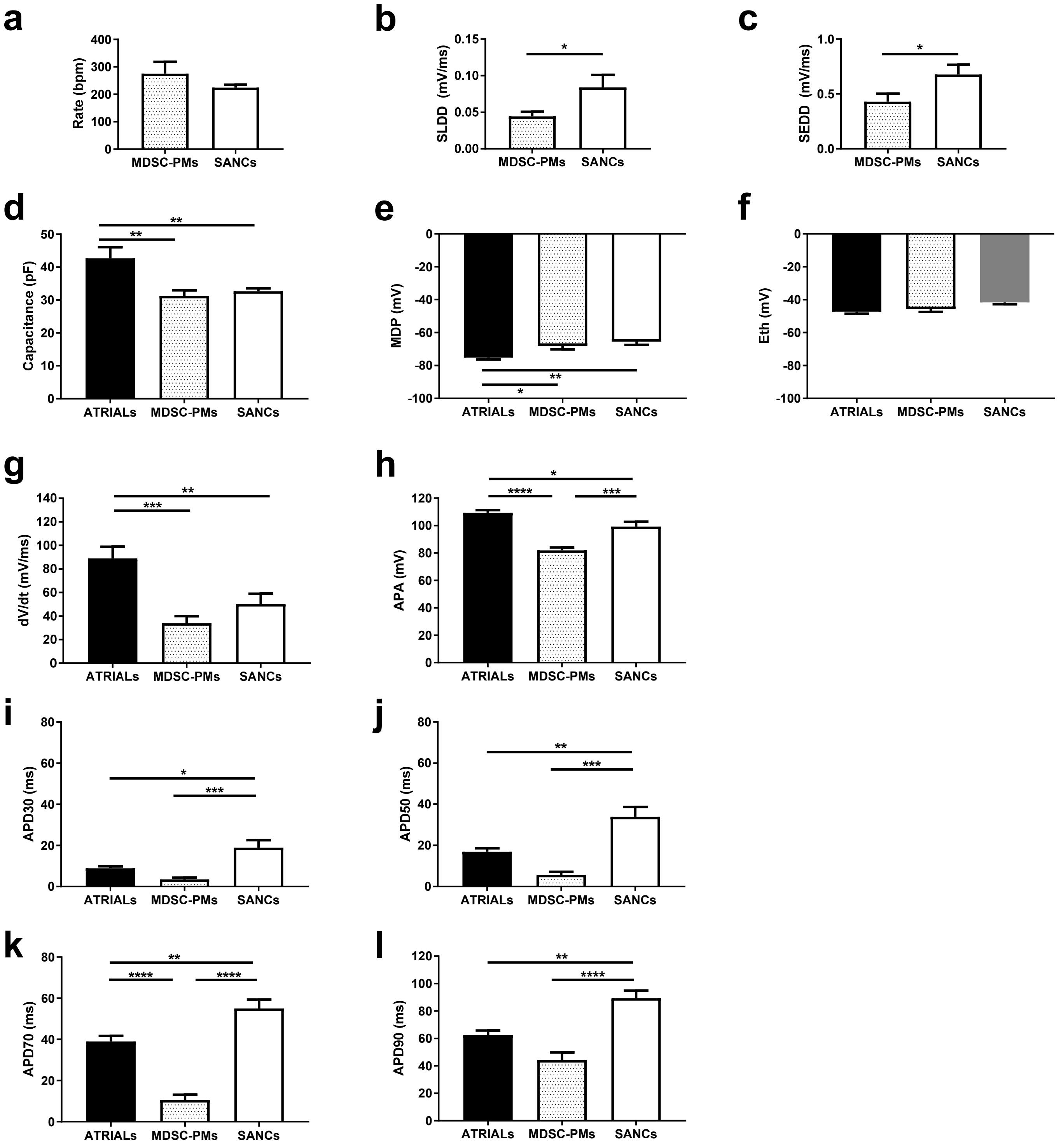


**Supplementary Fig. 3**. *Spontaneous action potential parameters of MDSC-PM and native SAN cells in comparison to evoked action potentials in atrial myocytes*. (**a**) spontaneous beating rate of MDSC-PM and native SAN pacemaker myocytes (SANCs); (**b**) slope of the linear phase of the diastolic depolarization (SLDD) and (**c**) slope of the exponential phase of the diastolic depolarization (SEDD); (**d**) membrane capacitance of MDSC-PM, native SANCs and atrial myocytes; (**e**) maximum diastolic potential (MDP); (**f**) threshold of the action potential ($E_{th}$); (**g**) first derivative of the action potential upstroke (dV/dt); (**h**) action potential amplitude (APA); (**i**-**l**) time at 30%, 50%, 70% and 90% of depolarization, respectively. Statistical significance was tested using unpaired t-test (a-c) and analysis of variance (ANOVA) followed by Tukey multiple comparisons test (d-i). *p<0.05, **p<0.01, ***p<0.001, ****p<0.0001. Error bars indicate s.e.m..

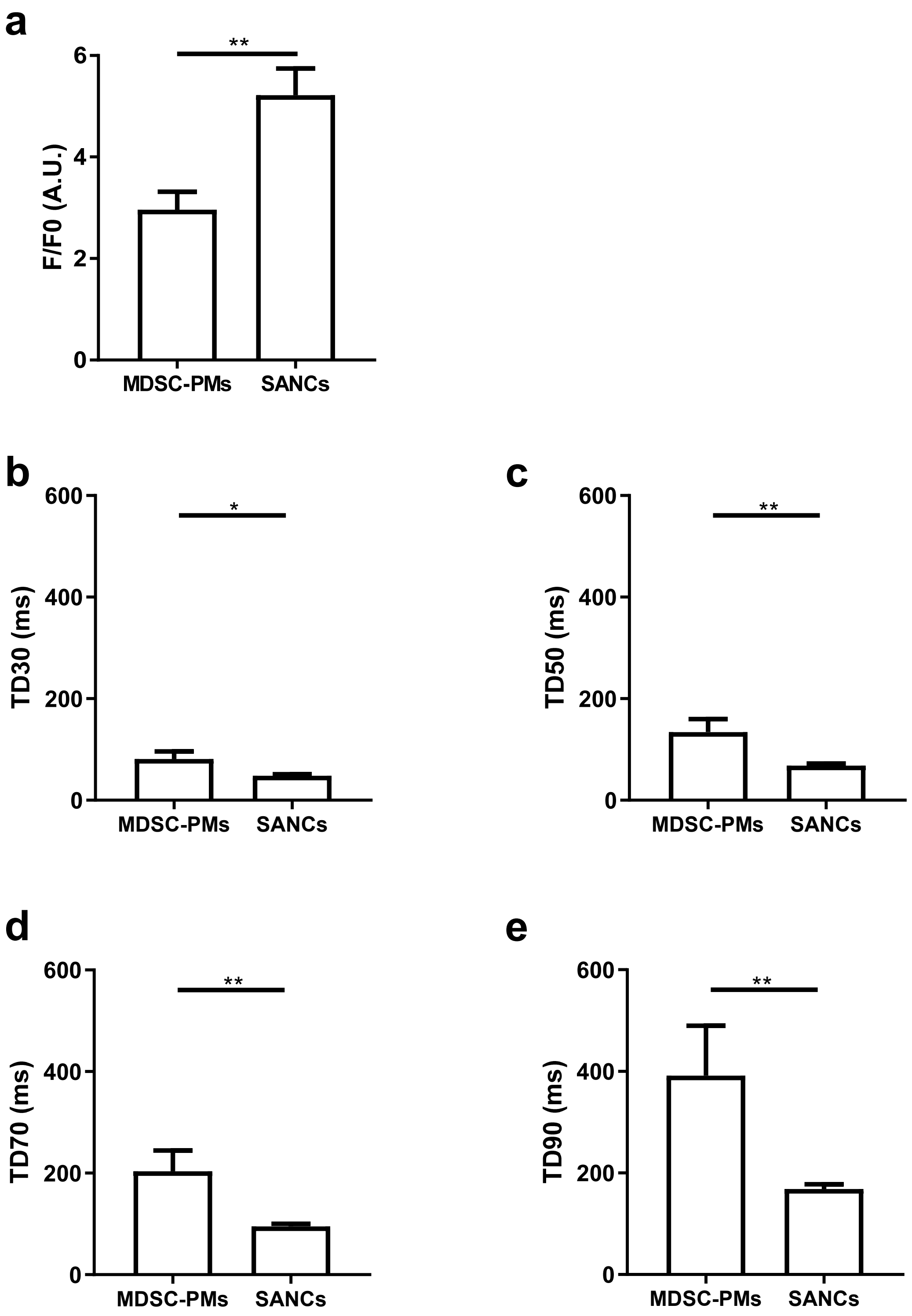


**Supplementary Fig. 4**. *Spontaneous $[Ca^{2+}]_i$ transients parameters of MDSC-PM and native SAN cells*. (**a**) Amplitude ($F/F_0$) of spontaneous $[Ca^{2+}]_i$ transients of MDSC-PM and native SAN pacemaker myocytes (SANCs); (**b**-**e**) Durations of spontaneous $[Ca^{2+}]_i$ transients at 30% (b), 50% (c), 70% (d) and 90% (e) of recovery. Statistical significance was tested using the unpaired t-test. *p<0.05, **p<0.01. Error bars indicate s.e.m..

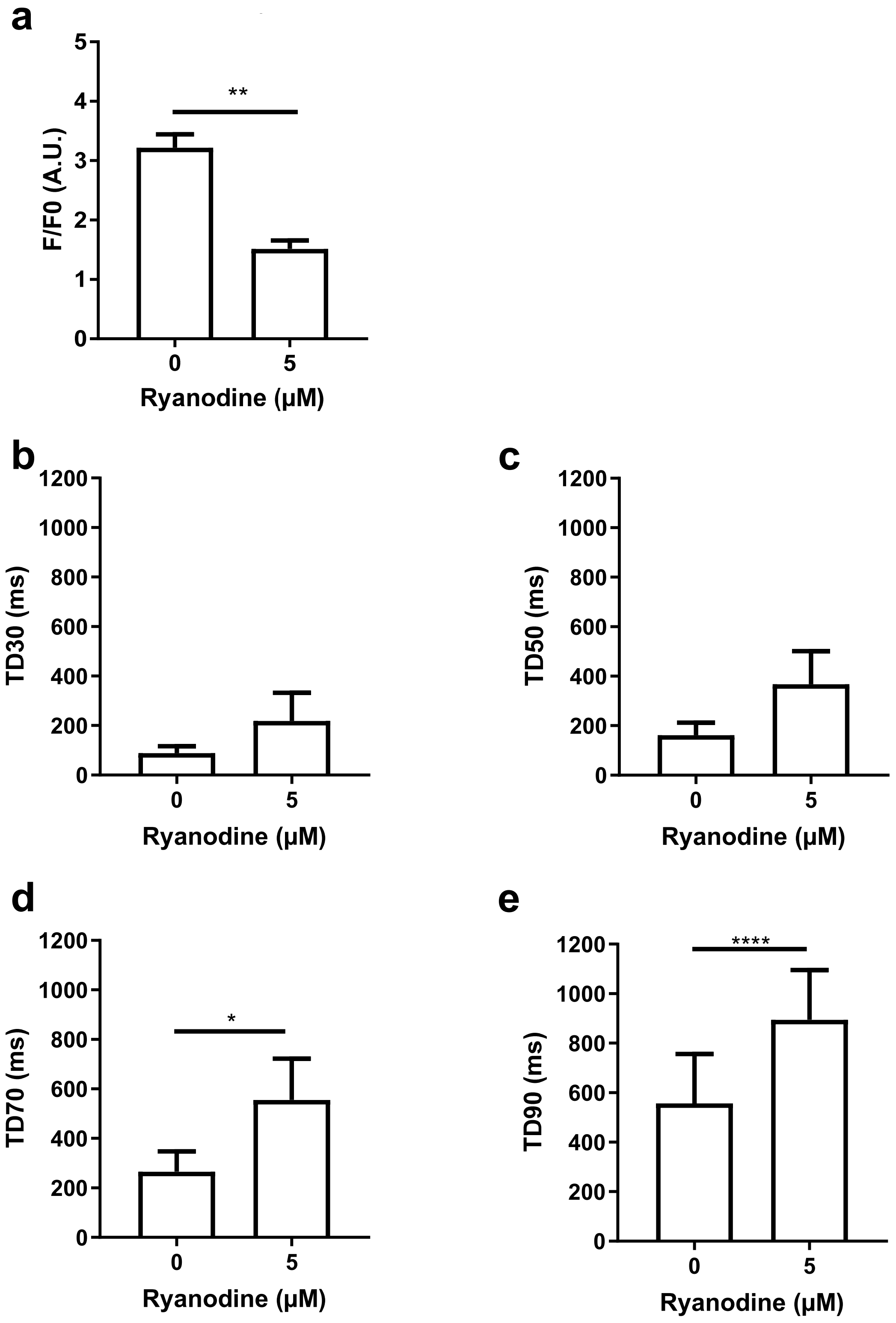


**Supplementary Fig. 5**. *Effects of ryanodine on spontaneous [$Ca^{2+}$]$_i$ transients in MDSC-PM*. (**a**) Reduction of the amplitude of spontaneous [$Ca^{2+}$]$_i$ transients by ryanodine. (**b**-**e**) Effects of ryanodine on the durations of spontaneous [$Ca^{2+}$]$_i$ transients at 30% (b), 50% (c), 70% (d) and 90% (e) of transient's recovery phase. Statistical significance was tested using the paired t-test. *p<0.05, **p<0.01. Error bars indicate s.e.m..

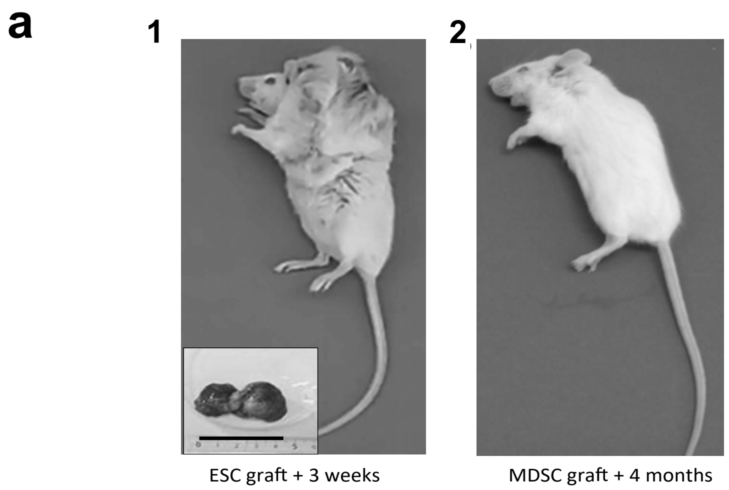


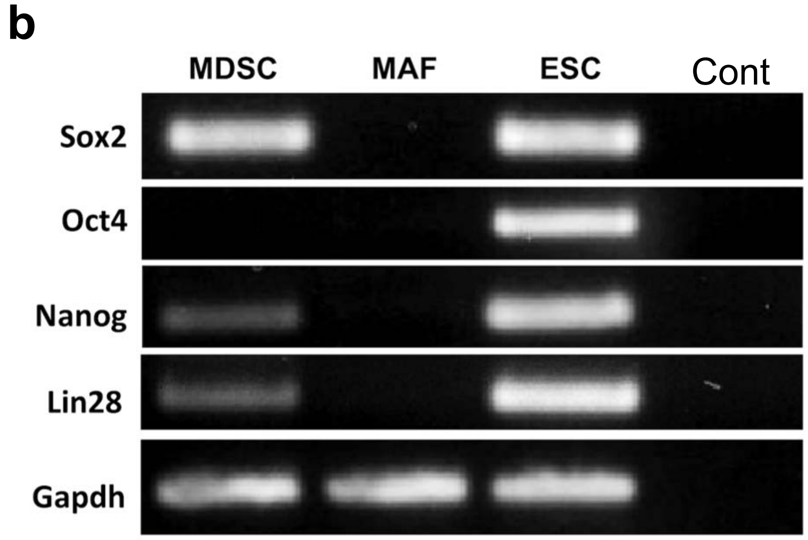


**Supplementary Fig. 6**. **a.** Tumor development following sub-cutaneous injection of mouse ESCs or MDSCs into SCID mice. 1. Sub-cutaneous injection of 1.106 mouse ESCs into SCID mice (n=2) induced fast-growing teratoma within 2 weeks. Inset shows the tumor extracted after 3 weeks; black scale bar = 4.5 cm. Panel 2. Sub-cutaneous injection of 1.106 MDSCs into SCID mice (n=4) failed to induce the formation of either teratomas or tumors for up to 4 months. **b.** RT-PCR-based expression analysis of MDSCs for transcription factors involved in stem-cell pluripotency: Sox2, Oct4, Nanog and Lin28. MDSCs: muscle-derived stem cells, MAF: mouse adult fibroblasts, ESCs: embryonic stem cells, Cont: control without cDNA, GAPDH was used as loading control

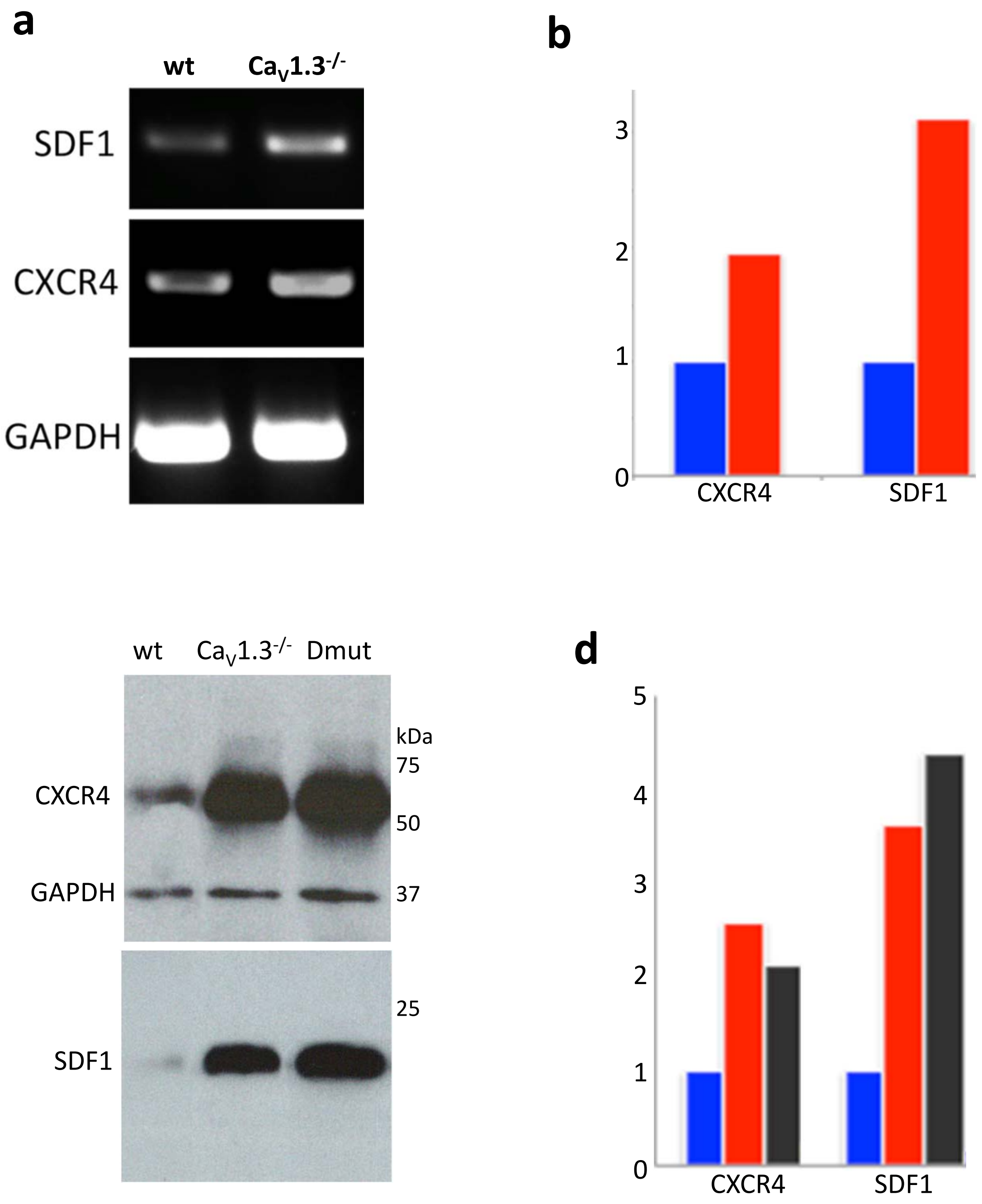


**Supplementary Fig. 7.** **(a)** RT-PCR-based analysis of expression of SDF1/CXCL12 chemokine and CXCR4 receptor in wild type(wt) and mutant $Ca_V1.3^{-/-}$ SAN plus RA tissue. Age- and sex-matched wild-type (wt) and $Ca_V1.3^{-/-}$ (mut) mice were used. GAPDH was used as loading control.
**(b)** Graphic representation of the quantitative analysis of the bands in SAN/RA tissue (a) blue bars in wild type samples, red bars in $Ca_V1.3^{-/-}$ mutant samples. **(c)** Western Blot analysis of CXCR4, GAPDH and SDF1 protein expression in wild-type, $Ca_V1.3^{-/-}$ mutant and $Ca_V1.3^{-/-}/Ca_V3.1^{-/-}$ double mutant (Dmut) in SAN tissue samples. **(d)** Graphic representation of the quantitative analysis of the bands in (c) normalized for GAPDH levels ) blue bars: wild type samples, red bars: $Ca_V1.3^{-/-}$ mutant samples, black bars: $Ca_V1.3^{-/-}/Ca_V3.1^{-/-}$ double mutant samples.

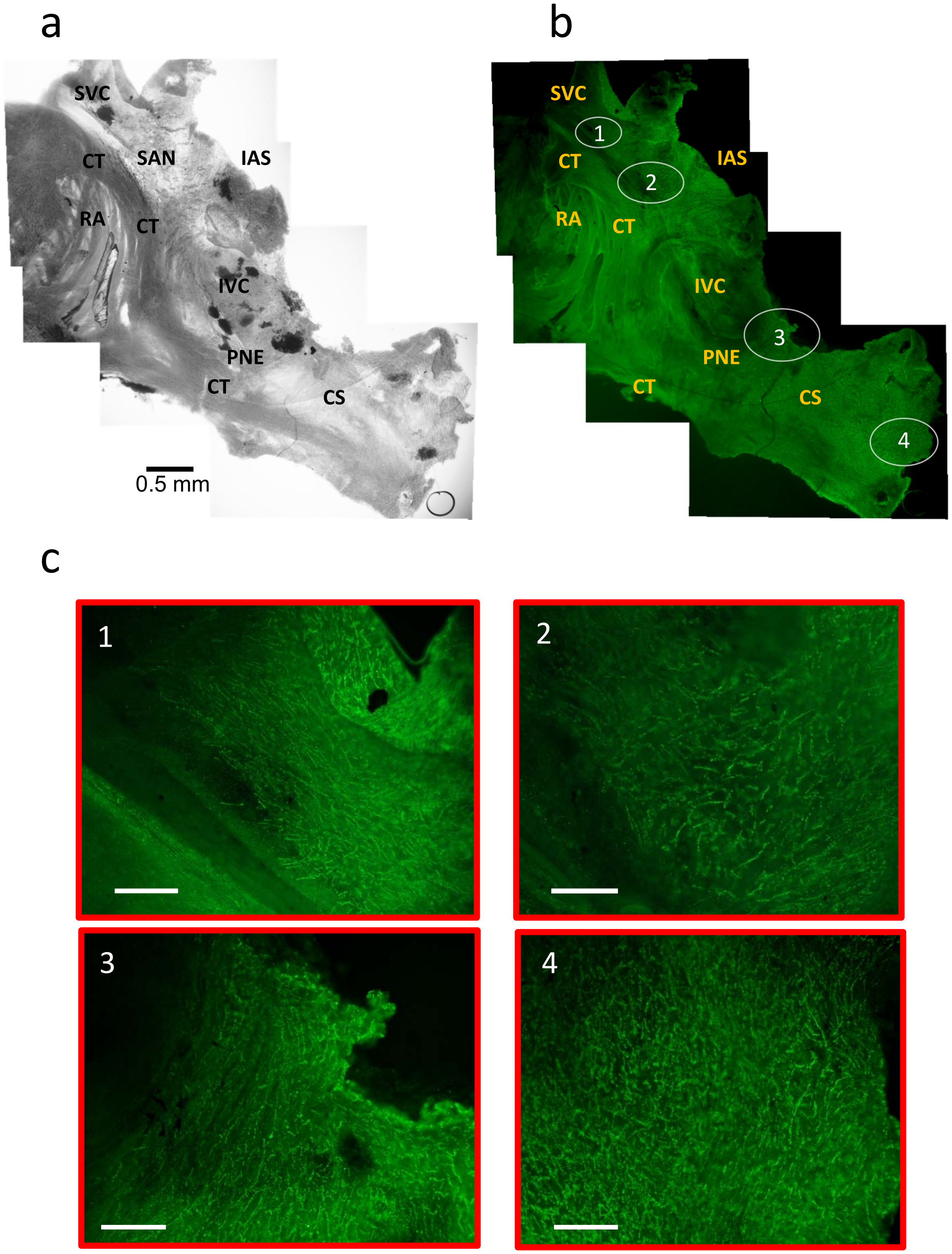
a
SVC
CT
SAN
IAS
RA
CT
IVC
PNE
CT
CS
0.5 mm
b
SVC
1
CT
2
IAS
RA
CT
IVC
3
PNE
CT
CS
4
c
1
2
3
4

(Bars: 100 μm)

**Supplementary Fig. 8.** (**a)** Phase-contrast image of isolated wild-type SAN. (**b**) Same tissue as in (a) immunolabelled with anti $Ca_v1.3$ antibody (see methods) showing the distribution of native expression of $Ca_v1.3$ channels. White circles 1-4 shows in the whole-mount SAN the position of close-ups in (c). (**c**) Close-up views of SAN corresponding to central areas (1 and 2), peripheral region corresponding to PNE (3) and the CS region (4). Abbreviations: SVC, superior vena cava; CT, crista terminalis; RA, right atrium; IAS, interatrial septum; IVC, inferior vena cava; CS, coronary sinus; PNE, posterior nodal extension.

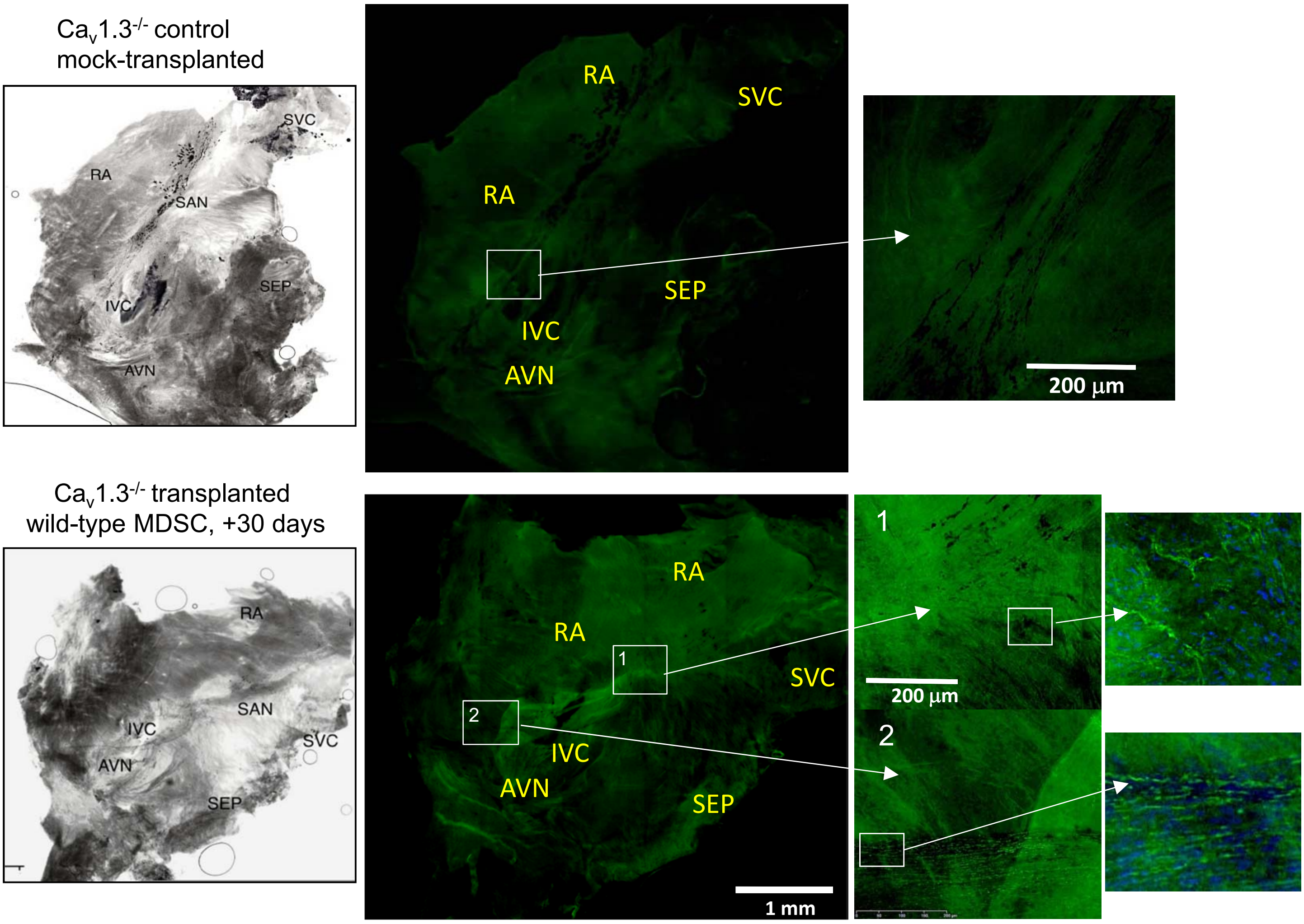


**Supplementary Fig. 9. Right atrial preparations containing the sino-atrial and atrioventricular nodes (SAN-AVN) from a mock-transplanted Ca$_V$1.3$^{-/-}$ mouse heart versus SAN-AVN from the heart of Ca$_V$1.3$^{-/-}$ mouse transplanted with wild-type MDSCs.**
SAN heart tissue from Ca$_V$1.3$^{-/-}$ mice was dissected 30 days after I.V. mock injection (0.9% NaCl, top panels) or I.V. MDSC injection (bottom panels) and immunostained for Ca$_V$1.3 as described in Fig. 6. Shown are whole mount SAN-AVN preparations scanned by Nanozoomer in brigthfield and green fluorescence, corresponding to Ca$_V$1.3 staining. Enlarged fields of areas with specific Ca$_V$1.3 staining of the transplanted SAN are shown in the right panels. Abbreviations: SAN, sino-atrial node; AVN, atrioventricular node; RA, right atrium; SVC, superior vena cava; IVC, inferior vena cava; SEP, inter atrial septum.

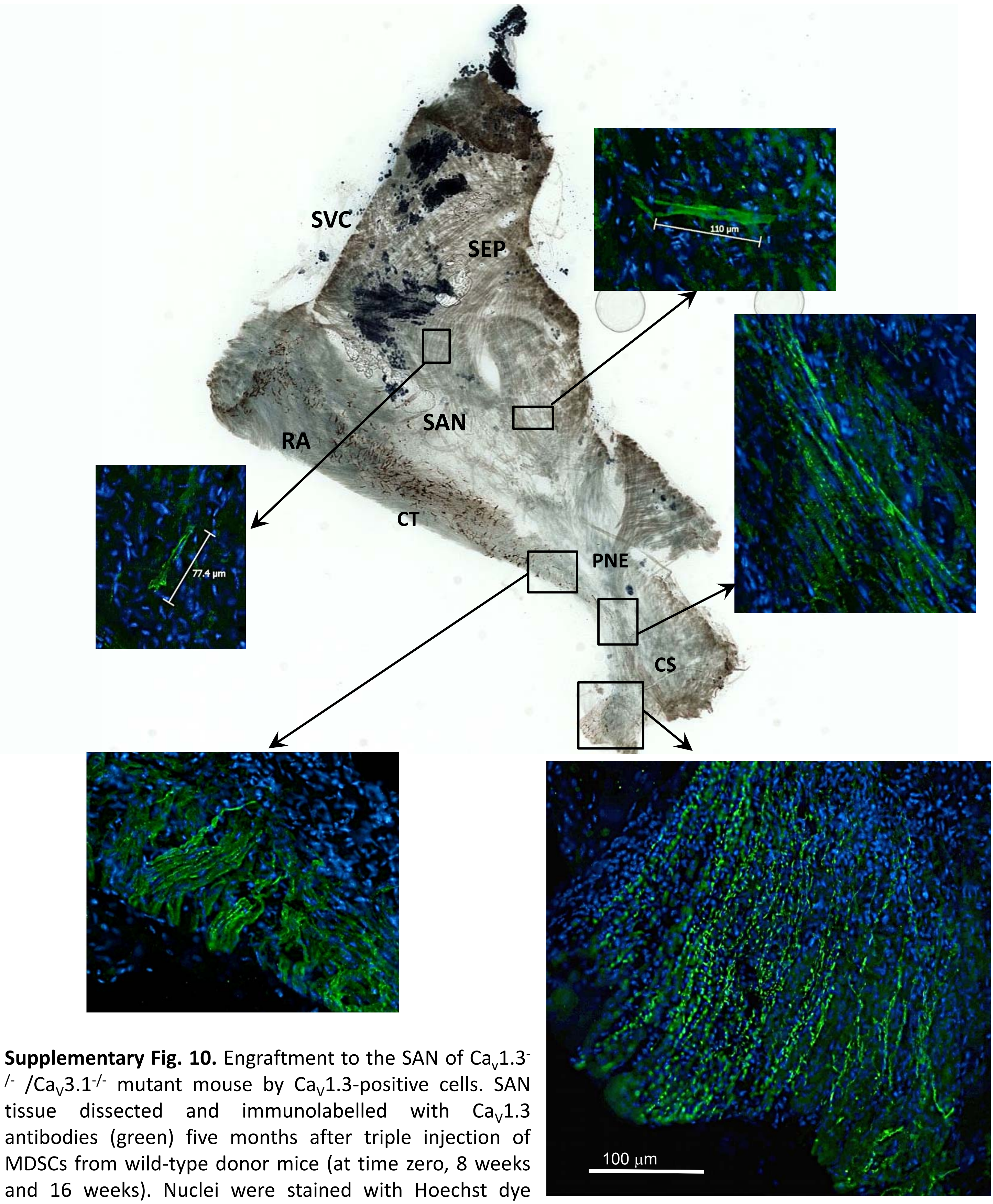


**Supplementary Fig. 10.** Engraftment to the SAN of $Ca_V1.3^{-/-}$ /$Ca_V3.1^{-/-}$ mutant mouse by $Ca_V1.3$-positive cells. SAN tissue dissected and immunolabelled with $Ca_V1.3$ antibodies (green) five months after triple injection of MDSCs from wild-type donor mice (at time zero, 8 weeks and 16 weeks). Nuclei were stained with Hoechst dye (blue). Large panel is bright-field image of SAN tissue. Enlarged panels show immunofluorescence signal for $Ca_V1.3$ and nuclei. Abbreviations: SAN, sino-atrial node; RA, right atrium; SVC, superior vena cava; SEP, inter atrial septum; CT, Crista Terminalis; PNE, Post Nodal Extension; CS, Coronary septum .

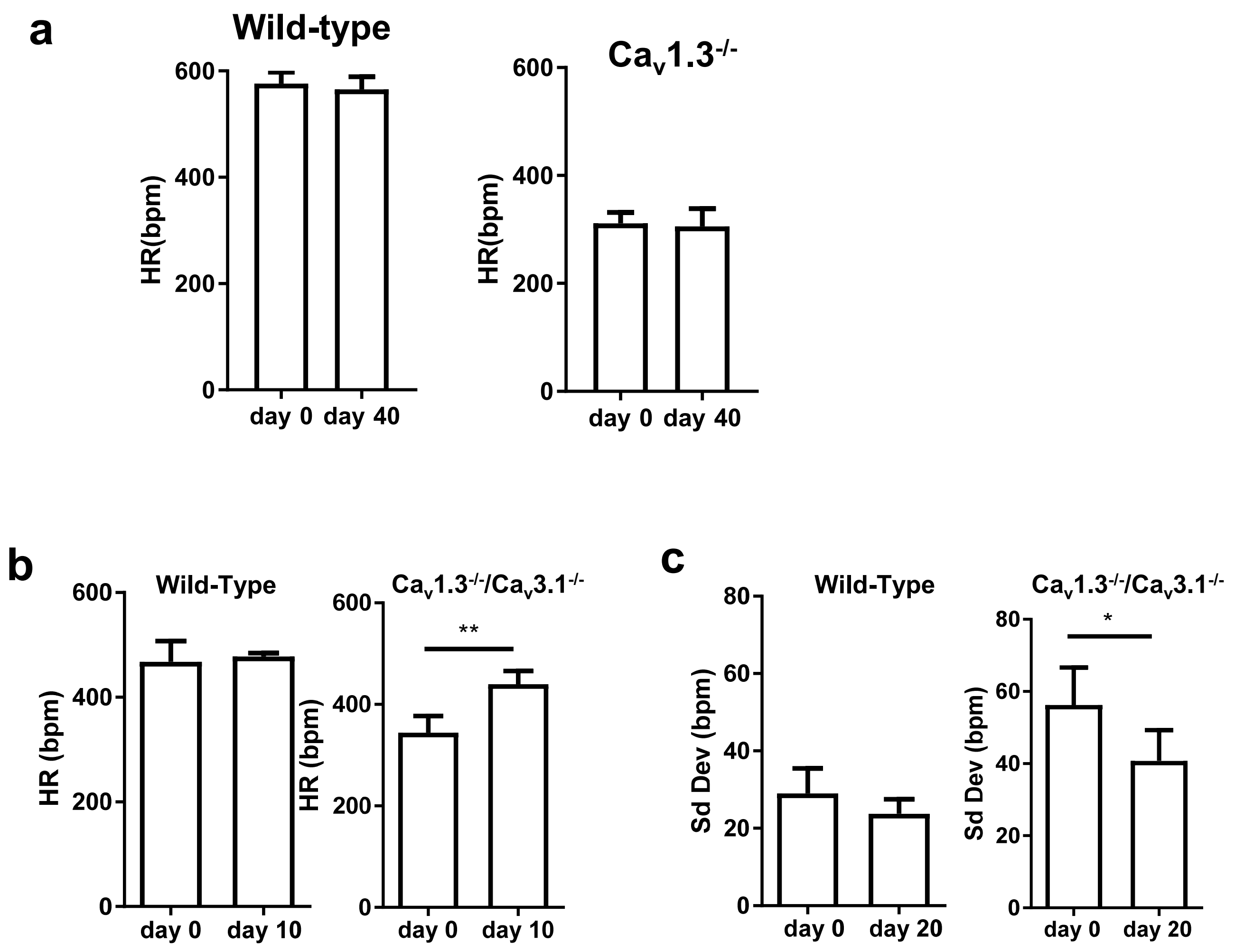


**Supplementary Fig. 11**. (**a**) **.** Bar graphs of heart rate (HR) after implant of a telemetric device and 40-days ECG follow-up recording in the absence of other treatment in age-matched wild-type (n=4) and $Ca_V1.3$-/- mice (n=5). Means are given ± S.E.M. (wild-type mice: 576 ± 21 to 565 ± 24 bpm; $Ca_V1.3$-/- mice: 311 ± 21 to 306 ± 29 bpm). (**b**). Bar graphs of the HR recorded in wild type (n=4) and $Ca_V1.3^{-/-}/Ca_V3.1^{-/-}$ (n=4) sedated mice before (day 0) and after (day 10) transplantation of MDSCs from wild-type mice. (wild type mice: 467 ± 40 at day 0 vs 478 ± 6 at day 10; $Ca_V1.3^{-/-}/Ca_V3.1^{-/-}$mice: 440 ± 26 at day 10, vs 344 ± 33 at day 0, p<0.01). (**c**) Standard deviation (Sd Dev) of HR recorded in control wild-type and $Ca_V1.3^{-/-}/Ca_V3.1^{-/-}$ mutant sedated mice before (day 0) and after (day 20) MDSC transplantation. The standard deviation of HR was significantly decreased 20 days after MDSC transplantation in $Ca_V1.3^{-/-}/Ca_V3.1^{-/-}$ mice. Statistical significance was tested using the paired t-test. *p<0.05, **p<0.01. Error bars indicate s.e.m..

**Supplementary Table 2. Telemetric ECG intervals for transplanted $Ca_v1.3^{-/-}/Ca_v3.1^{-/-}$ mice during 6 hours active (night) time.** Summary of ECG parameters recorded in n=6 $Ca_v1.3^{-/-}/Ca_v3.1^{-/-}$ double mutant mice before (day 0) and after (day 30) 30 days following I.P. transplantation of wild-type MDSCs. Shown are the values of the ECG parameters and the presence of sinus (SAN) pauses and 2nd –degree atrioventricular blocks (AVBII). Statistical significance was tested using the paired t-test. Data are presented as mean ± s.e.m..

| | Day 0 | Day 30 | n | p value |
|---|---|---|---|---|
| PP (ms) | 169 ± 8 | 135 ± 8 | 6 | 0.011 |
| PR (ms) | 54 ± 2 | 52 ± 1 | 6 | ns |
| QRS (ms) | 15 ± 1 | 15 ± 1 | 6 | ns |
| QT (ms) | 64 ± 3 | 62 ± 6 | 6 | ns |
| QTc (ms) | 50 ± 2 | 53 ± 4 | 6 | ns |
| SAN pauses/60s | 6 ± 2 | 2 ± 1 | 6 | ns |
| AVBII/60s | 6 ± 1 | 4 ± 2 | 6 | ns |
